# Coculture of *Collinsella aerofaciens* and *Bacteroides thetaiotaomicron* under bile acid stress reveals vitamin B6 exchange

**DOI:** 10.64898/2026.08.03.742426

**Authors:** Yan Wang, Sven-Bastiaan Haange, Andy Mercado Gamarra, Martin Leníček, Deborah Maier, Robin Plail, Hendrik Seidl, Christoph Kaleta, Ulrike Rolle-Kampczyk, Nico Jehmlich, Martin von Bergen

## Abstract

Although bile acid-mediated microbiome-host interactions are known to shape gut microbial composition and function, the mechanisms by which bile acid stress influences microbial metabolic interactions remain poorly understood. Here, we investigated the interaction between two key gut microbes, *Bacteroides thetaiotaomicron* and *Collinsella aerofaciens*, under deoxycholic acid (DCA) stress. In anaerobic coculture, *C. aerofaciens* mitigated the inhibitory effects of DCA on *B. thetaiotaomicron*, primarily through DCA uptake from the medium, as confirmed by DCA quantification. Proteomic analysis showed that DCA broadly disrupted amino acid and vitamin metabolism, particularly in *B. thetaiotaomicron*. In contrast, coculture promoted widespread metabolic activation in *C. aerofaciens*, including enhanced vitamin B6 metabolism and increased production of citrulline and ornithine. These findings suggest that metabolic cooperation enhances resistance to bile acid stress and may contribute to gut microbiome resilience, with potential relevance to liver- and bile acid-related disorders.

## Introduction

Bile acids are of particular interest in microbiome-host interaction. Bile acids make up approximately 80% of the contents of the gallbladder [1]. They act on bile acid receptors such as FXR and TGR5 and can regulate the composition of intestinal microbiota to influence metabolic health [2, 3]. The human liver synthesises two types of primary bile acids: cholic acid (CA) and chenodeoxycholic acid (CDCA). During digestion, primary bile acids are released into the small intestine to aid in the solubilization of lipids and lipid-soluble vitamins. Approximately 95% of primary bile acids are reabsorbed by the small intestinal wall and returned to the gallbladder, a process known as enterohepatic circulation. The remaining 5% enter the large intestine, where they come into direct contact with intestinal microorganisms and are converted into secondary bile acids, including deoxycholic acid (DCA) or lithocholic acid (LCA) [3–5].

On the side of the intestinal microbiome, bile acids influence microbial composition through their toxic effects on bacteria, including intracellular acidification, hydrogen sulfide release during taurine metabolism, and disruption of membrane integrity [4]. Dysregulation of bile acid metabolism can alter microbial composition and metabolic status and has been linked to liver diseases such as non-alcoholic fatty liver disease (NAFLD) and primary sclerosing cholangitis [6–8]

*Collinsella aerofaciens*, one of the most abundant members of the phylum Actinomycetota [9], is recognised as a bile acid-metabolising bacterium. It expresses bile salt hydrolase (BSH) and various hydroxysteroid dehydrogenases (HSDHs) that act on sterols, including bile acid 12α-HSDH [10, 11], as well as 7α- and 7β-HSDHs [11, 12], which are necessary for the production of hydrophilic secondary bile acids such as ursodeoxycholic acid (UDCA) [2, 13]. *C. aerofaciens* has also been reported to positively correlate with the production of DCA, particularly its glycine-conjugated form (gDCA) [2]. Despite its prevalence, the functional role of *Collinsella* remains largely underexplored. *C. aerofaciens* has been associated with various host conditions, including inflammatory bowel disease (IBD) [14, 15], irritable bowel syndrome (IBS) [16], NAFLD [17, 18] and low fibre diet [19]. These associations suggest that *C. aerofaciens* plays a complex and context-dependent role in microbiome-host interaction, which remains largely uncharacterised.

To better understand the functional role of *C. aerofaciens* within the gut microbiome, we established a defined coculture system with *Bacteroides thetaiotaomicron* in the presence of DCA. DCA is the single most abundant bile acid in the human gut [22]. Using integrated proteomic and metabolomic analyses, together with targeted quantification, we investigated how coculture shapes metabolic adaptation, with a particular focus on amino acid metabolism and vitamin biosynthesis. *Bacteroides thetaiotaomicron* is a prevalent (present in 46% of humans) [23] and abundant (up to 10^10^ per g stool) [24] gut symbiont widely used as a model organism for studying microbiome function. Together, this coculture system provides a tractable platform to dissect interspecies interactions relevant to bile acid stress and microbial contributions to gut–liver axis physiology.

## Methods

### Bacterial cultivation and bile acid stress assay

*Bacteroides thetaiotaomicron* DSM 2079 and *Collinsella aerofaciens* DSM 3979 were cultured anaerobically in BHIS medium at 37 °C and grown as mono- or cocultures in the presence or absence of 300 µM deoxycholic acid (DCA). Growth was monitored by OD_600_, and samples were separated into supernatant and cell pellet fractions for downstream analyses. Species abundances in cocultures were determined by species-specific absolute quantitative PCR (qPCR). Detailed procedures are provided in the Supplement.

### Metabolite analysis

Intracellular and extracellular DCA derivatives were analysed by LC–MS/MS as previously described [25]. Extracellular amino acids were quantified by PITC derivatisation followed by UPLC–MS/MS. Extracellular pyridoxal 5′-phosphate (PLP) was quantified using a fluorescence-based HPLC assay (KC2151, Immundiagnostik AG, Germany). Detailed procedures are provided in the Supplement.

### Metaproteome analysis

Proteins were extracted, digested by filter-aided sample preparation, and analysed by nanoLC–MS/MS on an Orbitrap Exploris 480 (Thermo Fisher Scientific) following Castañeda-Monsalve et al.[26]. Protein identification and quantification were performed in Proteome Discoverer (v2.5). Detailed protocols and instrument parameters are provided in the Supplement.

### Data analysis

Proteomics data quality control, statistical analyses and correlations were performed in R. Protein functions and pathway assignments were performed using Ghostkoala in KEGG database[27]. Functional analyses were conducted separately for *B. thetaiotaomicron* and *C. aerofaciens*. In the figures, they are abbreviated as B. theta / Bt and C. aero / Ca, respectively. Differential abundance analyses were performed using Kruskal–Wallis tests and two-way ANOVA. Metabolite concentrations and species abundances were analysed in GraphPad Prism 10. Detailed statistical procedures and significance criteria are described in the Supplement.

### Reference genome analysis

PLP biosynthesis and salvage capacities of *B. thetaiotaomicron* and *C. aerofaciens* were evaluated using an integrated workflow combining gapseq (v1.3)[28], KEGG orthology assignments (map ko00750)[27], and UniProtKB strain-level annotation[29]. Genome reconstructions were performed using the corresponding RefSeq assemblies[30]: *B. thetaiotaomicron* GCA_014131755.1 (ASM1413175v1) and *C. aerofaciens* GCF_010509075.1 (ASM1050907v1).

Gene presence was inferred from gapseq, KEGG, and UniProt annotations. Pathway completeness (de novo versus salvage) was inferred from combined annotations and compared with proteomic and targeted PLP data.

## Results

To investigate the impact of bile acid stress on microbial interactions, we compared *B. thetaiotaomicron* (*B. theta* / Bt) and *C. aerofaciens* (*C. aero* / Ca) in mono- and co-culture, under both control conditions and in the presence of DCA. Growth dynamics, proteomic profiles, and metabolite concentrations were measured to capture both physiological and metabolic changes.

### Coculture enhances tolerance to DCA

We first examined growth patterns under different culture conditions by measuring optical density at 600 nm (OD_600_) over time and assessing species abundance using quantitative PCR (qPCR). At the starting point (0 h) and after 3 h of incubation, no significant differences were observed among groups (Supplement Figure S1). After 6 h, however, DCA addition led to apparent growth inhibition in all cultures (Figure 1A). In *B. thetaiotaomicron* monoculture, OD_600_ values dropped by more than half (58%) in the presence of DCA, whereas *C. aerofaciens* showed only a modest decrease, indicating that *C. aerofaciens* is more tolerant of bile acid stress.

In coculture, the growth inhibition of *B. thetaiotaomicron* was substantially alleviated, as evidenced by an OD_600_ decrease of only 32% under DCA stress (Figure 1B). This suggests that interspecies interactions mitigate the inhibitory effect of bile acids on *B. thetaiotaomicron*. Interestingly, the opposite trend was observed for *C. aerofaciens* in coculture, where growth was somewhat reduced compared to its monoculture. However, growth was stronger in the presence of DCA, comparable to the abundance observed in monoculture with DCA. These results indicate a positive influence of *C. aerofaciens* on *B. thetaiotaomicron* in the presence of DCA, but an adverse effect of *B. thetaiotaomicron* on *C. aerofaciens* in coculture. It can be assumed that the competition for nutrients causes this, as *B. thetaiotaomicron* remained the dominant strain in coculture, supported by both 16S rRNA gene quantification and protein abundance (Figure 1C).

To further investigate the mechanism by which *C. aerofaciens* enhances the survival of *B. thetaiotaomicron* under bile acid stress, we quantified extracellular and intracellular concentrations of DCA and its derivatives (Figure 2). Compared to *B. thetaiotaomicron* monoculture, extracellular DCA levels were reduced in both *C. aerofaciens* monoculture and coculture, accompanied by an apparent increase in intracellular DCA within *C. aerofaciens*. In monoculture, the sum of extracellular and intracellular DCA totalled ∼400 µM, slightly above the nominal dose, likely due to analytical variation. Thus, these measurements cannot definitively confirm biotransformation. Nevertheless, the consistent total DCA across monoculture suggests that *C. aerofaciens* actively takes up DCA into its cells, lowering extracellular concentrations and thereby reducing *B. thetaiotaomicron*’s direct exposure to bile acid toxicity. Meanwhile, in the presence of *C. aerofaciens*, we detected the formation of glycine-conjugated DCA (Figure 2, right), although only at very low levels. In addition, minor amounts of 3′-keto-12α-hydroxy-DCA, 3α-hydroxy-12-keto-DCA, and taurine-conjugated DCA were observed. (Supplement Figure S2)

### Global proteomic and metabolite shifts

Proteomic analysis was performed to investigate the effects of bile acids on the two strains in mono- and co-culture. In total, 4,237 non-redundant proteins originating from *B. thetaiotaomicron* were identified, with an average of 3,694 ± 120 proteins detected per sample in those containing *B. thetaiotaomicron*. 2,537 proteins were identified from *C. aerofaciens*, with an average of 1,665 ± 375 proteins detected per sample in those containing *C. aerofaciens*.

Principal Component Analysis (PCA) revealed distinct effects of DCA on the proteomes of *B. thetaiotaomicron* and *C. aerofaciens.* In line with the growth results, DCA exerted a stronger impact on *B. thetaiotaomicron*, both in monoculture and coculture. For *B. thetaiotaomicron*, clusters with and without DCA treatment were clearly separated in monoculture, with no overlap, although greater variability was observed in the cocultured condition (Figure 3, left). In contrast, DCA had little apparent effect on *C. aerofaciens* (Figure 3, right). Instead, coculture was the primary factor shaping its protein expression. On the *B. thetaiotaomicron* side, coculture showed partial overlap of clusters with and without DCA, unlike in monoculture, indicating that *B. thetaiotaomicron* functioned more similarly under the influence of *C. aerofaciens.* Notably, *B. thetaiotaomicron* monoculture under DCA stress was distinct from *B. thetaiotaomicron* in coculture under DCA stress, supporting the hypothesis that *C. aerofaciens* alleviates DCA-induced stress in coculture (Figure 3, left). *C. aerofaciens* clusters from coculture with DCA overlapped partially with monoculture clusters regardless of DCA treatment, whereas coculture without DCA formed distinct clusters, suggesting potential cross-feeding interactions in which *C. aerofaciens* benefited from *B. thetaiotaomicron* when DCA did not impair the latter (Figure 3, right).

The more detailed analysis revealed pathway-specific effects of DCA treatment and of coculture (Figure 4). Focusing on the effect of DCA, *B. thetaiotaomicron* as a whole showed a suppressed state, with most of them appearing down-regulated compared to the absence of DCA stress, except for selenocompound metabolism. *C. aerofaciens*, on the other hand, did not show significant changes in stable occurrence in monoculture and coculture (Figure 4, left), which also contributes to the finding that *C. aerofaciens* was not as affected as *B. thetaiotaomicron* by DCA. The alterations caused by coculture (Figure 4, right) revealed that in *B. thetaiotaomicron*, the pathways for valine, leucine, and isoleucine (branched-chain amino acids, BCAAs) biosynthesis, as well as cysteine and methionine metabolism, were upregulated, accompanied by an increase in pantothenate (vitamin B5, VB5) levels. In *C. aerofaciens*, metabolic alterations were more pronounced, including upregulation of cysteine and methionine metabolism, lysine metabolism, pantothenate and CoA biosynthesis, selenocompound metabolism, and, notably, vitamin B6 metabolism. These pathways indicate potential cross-feeding or stress responses in the presence of *B. thetaiotaomicron*.

To evaluate whether some of the produced amino acids contribute to metabolic cross-feeding, the bacterial culture supernatants were analysed by targeted metabolomics, focusing on amino acids and biogenic amines. Under DCA treatment, larger effect sizes were associated with higher amino acid concentrations in the DCA-treated groups (Figure 5, left). This may reflect suppression of metabolic pathways and reduced biomass due to growth inhibition, or alternatively, active production of specific metabolites. The impact was more pronounced in coculture (Figure 5, right). Notably, citrulline concentrations were significantly higher in coculture compared with *B. thetaiotaomicron* monoculture. Ornithine and spermidine levels also increased in coculture, but only in the absence of DCA. This suggests an influence of *C. aerofaciens.* When comparing coculture with *C. aerofaciens* monoculture, increases in more amino acids were observed, including glutamate, alanine, BCAAs, and, in particular, aspartate, glycine, and serotonin. *B. thetaiotaomicron* likely contributes these metabolites.

### Vitamin B6 biosynthesis is enhanced in coculture

Vitamin B6 in its active form, pyridoxal 5’-phosphate (PLP), serves as a versatile cofactor in numerous enzymatic reactions, including those central to amino acid metabolism and cellular stress protection. In our global proteomic profiling, we observed a notable enrichment of PLP-related enzymes in coculture compared to monoculture (Figure 4, right). Targeted PLP quantification assay revealed that coculture of *B. thetaiotaomicron* and *C. aerofaciens* led to a pronounced enhancement of vitamin B6 biosynthesis (Figure 6). In monoculture, *B. thetaiotaomicron* primarily acted as a consumer of PLP, resulting in reduced extracellular PLP levels, while *C. aerofaciens* alone showed no appreciable effect on PLP concentrations. By contrast, in coculture, we detected a significant accumulation of extracellular PLP. Proteomic data further revealed that the PLP synthase subunits pdxS and pdxT were strongly upregulated in *C. aerofaciens* under coculture conditions, in parallel with the increase in metabolites (Figure 6).

This coordinated pattern suggests that the presence of *B. thetaiotaomicron* stimulates *C. aerofaciens* to activate PLP biosynthesis, thereby enhancing extracellular vitamin B6 availability.

To directly assess whether vitamin B6 availability could benefit *B. thetaiotaomicron*, the impact of supplementation with defined PLP concentrations was analysed (Supplement Figure S3). At low concentrations (≤50 ng/L), PLP supplementation promoted *B. thetaiotaomicron* growth, with significant effects observed from the 6th hour of culture at a concentration of 50 ng/L. A smaller but consistent, though not statistically significant, growth-promoting trend was also observed at 10 ng/L. By contrast, supplementation with higher PLP concentrations (≥100 ng/L) inhibited growth, possibly reflecting cytotoxic effects. These results confirm that vitamin B6 availability can modulate *B. thetaiotaomicron* growth dynamics in a dose-dependent manner, and the concentrations observed in coculture fall within the growth-promoting range.

### PLP metabolism pathway completeness differs between strains

To explore whether intrinsic biosynthetic differences might underlie the vitamin B6 patterns observed experimentally, the reference genomes of *B. thetaiotaomicron* DSM 2079 and *C. aerofaciens* DSM 3976. Annotation across gapseq, KEGG, and UniProt suggested differences in pathway completeness between the two strains.

In *B. thetaiotaomicron*, enzymes such as pdxA, serC, and pdxH were supported by the consulted annotation sources, indicating that upstream precursor formation and vitamin B6 interconversion reactions are likely encoded. However, the core PLP synthase subunits pdxS and pdxT lacked support in both gapseq (“bad blast” or “no blast”) and UniProt (no *B. thetaiotaomicron*-specific accessions). Additionally, other de novo-associated enzymes (e.g., pdxJ) were weakly supported. These findings suggest that *B. thetaiotaomicron* DSM 2079 may lack a complete de novo PLP biosynthetic pathway, although salvage and interconversion reactions are likely present.

In *C. aerofaciens*, evidence for the pathway was more complete. For *C. aerofaciens*, none of the essential PLP biosynthesis genes were unsupported across all tools. At least one annotation system (gapseq BLAST scores, KEGG KO mapping, or UniProtKB strain-level protein IDs) indicated the presence of pdxS, pdxT, pdxB, pdxJ, serC, and pdxK. For example, gapseq identified pdxS with a “good blast hit”, pdxT with a lower-confidence hit, and UniProt returned *C. aerofaciens*-specific accessions for both subunits. KEGG also mapped these enzymes to the strain. Although the strength of support varied, each of these genes had at least partial independent evidence for presence, which may indicate that *C. aerofaciens* DSM 3979 could encode a functional de novo PLP pathway.

Overall, the genomic evidence supports a model in which *C. aerofaciens* would be PLP-proficient, whereas *B. thetaiotaomicron* would be PLP-dependent. These differences provide a genomic basis for the species-specific proteomic and metabolic responses observed in mono- and coculture.

### Arginine metabolism shifts toward stress-protective pathways

Arginine-related metabolic pathways in *C. aerofaciens* displayed notable shifts during coculture. These changes were evident at both the metabolomic and proteomic levels. Proteomic profiling (Figure 4, right) revealed an upward trend in arginine biosynthesis in coculture, which was particularly pronounced under DCA stress. Comparison of *C. aerofaciens* monoculture and coculture further showed a larger effect size in the latter (Figure 5, right), suggesting that *B. thetaiotaomicron* influenced arginine metabolism in *C. aerofaciens*.

Metabolomic analyses from the supernatant (Figure 7) confirmed alterations in arginine and related intermediates. While arginine, ornithine, and citrulline were overall consumed, their levels differed between species. In *B. thetaiotaomicron* monoculture, arginine concentrations remained relatively higher, whereas in *C. aerofaciens* monoculture, ornithine and citrulline levels were elevated compared to the other groups. These patterns suggest species-specific compensatory production of intermediates within the arginine pathway. These observations are consistent with active conversion of arginine into ornithine and citrulline by *C. aerofaciens*, with citrulline accumulation particularly enhanced under DCA stress. This interpretation is supported by proteomic evidence showing upregulation of enzymes initiating the unidirectional conversion of glutamate to ornithine, thereby feeding into the arginine–citrulline–ornithine cycle. Additionally, these shifts may extend to polyamine metabolism, as spermidine (SPD) displayed a similar accumulation pattern. Given the established role of spermidine in bacterial stress tolerance, this observation may indicate a link between arginine metabolism and adaptation to bile acid stress.

### Interaction analysis identifies pathways protected in coculture

To determine whether DCA exerted a stronger effect in coculture, limma, a two-way ANOVA method, was applied (Figure 8). The analysis showed that arginine metabolism, cysteine and methionine metabolism, and selenocompound metabolism differed significantly between coculture and monoculture in both strains.

## Discussion

### Bile acid stress and microbial redox balance

Bile acids such as DCA impose oxidative stress on gut bacteria through their hydrophobicity, disrupting membrane integrity and causing redox imbalance and cellular damage [3]. In this study, *C. aerofaciens* was found to partially alleviate this stress, primarily through bile acid uptake and, to a lesser extent, modification. Supporting this, small amounts of glycine-conjugated DCA (gDCA) were detected in both culture supernatants and cell pellets (Figure 2). However, *C. aerofaciens* does not appear to be a major contributor to bile acid conjugation. Consistent with this observation, proteomics detected no enzymes involved in DCA biotransformation. Bacteria can sequester drugs intracellularly without chemical modification, a process known as bioaccumulation. Although this may not affect growth, intracellular sequestration can perturb cellular metabolism and promote cross-feeding interactions within microbial communities [31].

While gDCA is the predominant conjugated form in humans [2], trace levels of taurine-conjugated DCA, commonly found in rodents, and other optimised DCA derivatives were also detected, albeit at very low abundance (Supplement Figure S2). Although DCA concentrations decreased in supernatants following *C. aerofaciens* uptake in both mono- and cocultures, with the greatest decrease in coculture, accumulation in cell pellets was not proportional, and only trace amounts were detected (Figure 2). This discrepancy may reflect (i) technical limitations in cell disruption and metabolite extraction, particularly for *C. aerofaciens*, a bacterium involved in lipid metabolism [18], where extraction efficiency may differ; or (ii) the possibility that *C. aerofaciens* and *B. thetaiotaomicron* synergistically generate unconventional bile acid conjugates not captured by the analytical approach used. Indeed, conjugation has been reported not only with glycine and taurine but also with alanine, proline, leucine, and phenylalanine [32], as well as fatty acids [33]. Furthermore, cysteamine-conjugated bile acids have recently been identified in a murine knockout model deficient in glycine/taurine conjugation [34]. These findings highlight the expanding diversity of bile acid conjugation chemistry and suggest bacterial contributions may extend beyond classical glycine and taurine conjugates.

The synergistic interaction between *C. aerofaciens* and *B. thetaiotaomicron* in coculture may further mitigate DCA-induced oxidative stress. Widespread changes in protein expression were observed in response to DCA, with cocultures exhibiting distinct profiles from monocultures (Figure 3 & Figure 4). Pathways related to amino acid metabolism, vitamin biosynthesis, and stress defence were particularly of interest (Figure 4 & Figure 5). Targeted metabolite analysis confirmed alterations in extracellular amino acids, vitamins, and bile acid derivatives, indicating coordinated metabolic adaptation between the two species. Notably, although DCA broadly suppressed metabolic pathways in *B. thetaiotaomicron* monoculture, selenocompound metabolism increased in abundance (Figure 4, left), suggesting activation of a stress-responsive defence mechanism.

Limma analysis further revealed positive interaction effects for these pathways (Figure 8). Given that arginine and cysteine/methionine metabolism were suppressed by DCA, while selenocompound metabolism showed a strain-dependent response (Figure 4, left), the positive interaction terms indicate that this suppression was attenuated in coculture and, in some cases, reversed. Interaction plots showed that cysteine and methionine metabolism was suppressed by DCA in both strains, regardless of culture condition, whereas arginine and selenocompound metabolism in *C. aerofaciens* displayed opposite trends in monoculture and coculture (Figure 8, B). Figure 8, therefore, identifies arginine, cysteine, and selenocompound metabolism as pathways whose inhibition by DCA was attenuated in coculture.

These pathways are closely linked to detoxification. Arginine metabolism feeds into ornithine and subsequently polyamine synthesis, and polyamines are known antioxidants [35]. Cysteine, a sulfur-containing amino acid, provides reducing power and serves as a precursor for glutathione [36]. Selenocompounds contribute through the formation of selenoproteins such as glutathione peroxidases and can directly bind and facilitate the excretion of heavy metals [37].

### Vitamin B6 Biosynthesis as a Stress-Responsive Pathway

As the active form of vitamin B6, PLP serves as a cofactor in numerous metabolic reactions in both hosts and microorganisms. Beyond its role in amino acid metabolism, PLP has been linked to microbial oxidative-stress responses and redox homeostasis. The increased PLP abundance observed in coculture therefore suggests that vitamin B6 metabolism may contribute to adaptation under DCA exposure.

Although *C. aerofaciens* has been proposed to synthesise vitamin B6 [38], this is currently supported only by genomic predictions and lacks experimental validation. In this study, coculture was associated with enhanced PLP synthesis. Although the increase may primarily originate from *C. aerofaciens*, *B. thetaiotaomicron* could also contribute through vitamin B6–PLP interconversion. Genomic analyses suggest that Bacteroides species may participate via pathways involving deoxy-xylulose 5-phosphate and 4-phosphohydroxy-L-threonine [38]. Although *B. thetaiotaomicron* is not generally considered capable of de novo PLP biosynthesis, it possesses enzymes that convert other vitamin B6 vitamers into PLP. Thus, alongside production by *C. aerofaciens*, *B. thetaiotaomicron* may contribute through vitamin B6–PLP interconversion, supporting its own growth and metabolism.

To explore potential mechanisms underlying the enhanced PLP production, we next examined metabolites and proteins associated. In our metabolomic analysis, aspartate and asparagine displayed distinct patterns between monoculture and coculture (Figure 5, right). While *C. aerofaciens* rapidly depleted available Asp in monoculture, *B. thetaiotaomicron* also consumed Asp but consistently maintained residual levels in the medium (Figure 9A). Notably, in coculture, *B. thetaiotaomicron* cell numbers remained similar (Figure 1B), yet overall Asp depletion was not substantially greater despite the presence of the additional consumer, *C. aerofaciens* (Figure 9A). These observations are consistent with the possibility that *B. thetaiotaomicron* partially compensates for the increased demand by supplying Asp during coculture. Proteomic data are consistent with this interpretation, as *B. thetaiotaomicron* upregulated K01424, an enzyme converting asparagine to aspartate. Meanwhile, *C. aerofaciens* induced the aspartate transporter (K11358) together with enzymes involved in glutamine biosynthesis (Figure 9A). As glutamine serves as a key precursor for de novo PLP synthesis, these changes are consistent with increased flux towards vitamin B6 production. Although aspartate aminotransferase was detected inconsistently, its high abundance when present suggests a potentially important role in directing aspartate flux towards PLP synthesis. Collectively, these findings support a model in which *B. thetaiotaomicron* supplies nitrogenous precursors, whereas *C. aerofaciens* converts them into glutamine and subsequently PLP, resulting in a coordinated enhancement of vitamin B6 metabolism during coculture.

Aminotransferases constitute the largest group of PLP-dependent enzymes [39], and their activities are central to amino acid metabolism and microbial cross-feeding. In coculture, upregulation of several PLP-dependent enzymes suggests that increased PLP availability supports broader metabolic functions (Figure 9B & Supplement Figure S4). In *C. aerofaciens*, increased expression of diaminopimelate decarboxylase, O-acetylhomoserine thiol-lyase, and cysteine synthase points to enhanced lysine, cysteine and methionine synthesis, with the latter two enzymes increasing more than twofold (Figure 9B). In *B. thetaiotaomicron*, upregulated enzymes included BCAA aminotransferase, reinforcing its role in BCAA turnover, and aspartate 4-decarboxylase, linking aspartate utilisation to β-alanine and pantothenate (vitamin B5) biosynthesis. Together, these changes suggest that enhanced PLP availability promoted not only amino acid interconversion within each species but also complementary metabolic exchanges between them, particularly within sulfur amino acid metabolism.

Importantly, enhanced cysteine metabolism in *C. aerofaciens* may contribute to alleviating DCA-induced oxidative stress. Consistent with this interpretation, DCA-induced suppression of cysteine and methionine metabolism was attenuated in coculture, together with increased abundance of PLP-dependent enzymes involved in these pathways. This interpretation is consistent with the broader role of vitamin B6 in redox homeostasis. Although less recognised than vitamins C or E for direct antioxidant activity [40], vitamin B6 indirectly supports antioxidant defence by promoting cysteine synthesis, thereby fuelling glutathione production and the transsulfuration pathway [41, 42]. In addition, pyridoxine and pyridoxamine possess direct radical-scavenging activity and can quench reactive oxygen species (ROS) [40]. Thus, enhanced vitamin B6 production in coculture may provide dual protection against oxidative stress by reinforcing sulfur-dependent redox metabolism while also supplying direct antioxidant capacity. Beyond microbial stress adaptation, enhanced vitamin B6 production may also be relevant to host gut–liver axis health [43–46].

### Metabolic Cross-Feeding Between *B. thetaiotaomicron* and *C. aerofaciens*

Metabolic cross-feeding emerged as a central feature of the coculture between *B. thetaiotaomicron* and *C. aerofaciens* (Figure 10). Proteomic and metabolite data indicated that *C. aerofaciens* metabolism was broadly stimulated in coculture, likely driven by nutrient inputs from *B. thetaiotaomicron.* In particular, arginine utilisation enabled downstream ornithine and citrulline production, while BCAA metabolism showed evidence of cross-feeding, consistent with the upregulation of *C. aerofaciens* BCAA transporters (Supplement Figure S5). Enhanced pantothenate synthesis in *B. thetaiotaomicron* may further contribute to metabolic stability within the consortium. *C. aerofaciens* consistently produced ornithine and citrulline, with citrulline accumulating to higher concentrations. These patterns are consistent with the activity of the arginine deiminase (ADI) pathway, which generates ATP and ammonium from arginine metabolism. Under hypoxic conditions, this pathway may support energy production, while ammonium release could help buffer acidification caused by acetate production by *B. thetaiotaomicron* output (Supplement Figure S6) [47]. Spermidine accumulation followed a similar trend to ornithine and citrulline, suggesting that arginine metabolism may extend into polyamine production. As spermidine is associated with bacterial stress tolerance, this pathway may contribute to adaptation under bile acid stress. Although genomic annotations suggest *C. aerofaciens* lacks a complete spermidine biosynthetic pathway, it remains possible that relevant enzymes are present but not yet characterised, leaving its precise origin unresolved.

Evidence for BCAA cross-feeding was also observed. *B. thetaiotaomicron* is a recognised degrader of BCAAs and functions as a metabolic sink [8], whereas *C. aerofaciens* upregulated BCAA transporters in coculture. Although the physiological significance of this interaction remains unclear, it may represent a previously unrecognised metabolic link between the two species. Given the association between altered BCAA metabolism and NAFLD [48].

### Limitations and Future Work

This study employed *B. thetaiotaomicron* and *C. aerofaciens* to analyse the effects of bile acid on the physiology of two essential species from the human microbiome and their potential metabolic interaction. By combining coculture under DCA stress with proteomic profiling and targeted metabolite quantification, we captured key aspects of their functional dynamics. Nonetheless, microorganisms rarely exist in isolation; instead, they form densely interconnected communities through nutrient exchange and the production of metabolic by-products. Our two-species coculture represents a highly simplified system that cannot fully reflect the complexity, stability, or continuous nutrient environment of the gut. Moreover, bile acid concentrations and compositions vary considerably in vivo, and thus the conditions tested here only approximate the physiological range.

From a methodological perspective, metabolomics provided only a limited snapshot, lacking insights into ongoing metabolic fluxes. Additionally, disentangling the precise contributions of each species to shared metabolites remains challenging. Addressing these limitations will require isotope tracing to resolve metabolic fluxes, as well as validation in more complex and physiologically relevant systems, such as multi-species consortia, continuous bioreactors, or host–microbe models.

Microbial interactions extend beyond cross-feeding and include both exploitative and interference competition strategies [49]. In this study, we were unable to explore these dynamics further. Likewise, analyses can extend beyond transporter profiling to include quorum-sensing (QS) systems [50], which may contribute to the regulation of amino acid exchange, PLP biosynthesis, and arginine metabolism in coculture.

Physical interactions between bacteria constitute another underexplored dimension of interspecies dynamics. Direct cell–cell contact can facilitate nutrient exchange, chemical signalling, and horizontal gene transfer [51], while bacterial aggregates and biofilm-like structures may create localised microenvironments that alter pH, redox potential, and bile acid concentrations [52]. In our study, we did not determine whether *B. thetaiotaomicron* and *C. aerofaciens* form aggregates or exhibit direct adhesion under bile acid stress. Future microscopy-based approaches combined with cell-labelling could reveal co-localisation patterns, while single-cell techniques may further elucidate metabolic coupling within aggregates.

Looking forward, the insights gained here highlight how *C. aerofaciens* contributes to maintaining redox balance and facilitating nutrient interconversion under bile acid stress, suggesting potential roles in shaping host–microbe interactions in both health and disease. Future studies extending this work into complex communities and in vivo models may clarify whether modulating *C. aerofaciens* activity can be leveraged to counteract oxidative stress, improve bile acid tolerance, and mitigate progression of liver-related disorders such as NAFLD.

## Conclusions

This study demonstrates that coculture of *B. thetaiotaomicron* and *C. aerofaciens* promotes adaptation to DCA stress through coordinated metabolic responses. *C. aerofaciens* contributed to bile acid uptake, while coculture enhanced vitamin B6 metabolism and alleviated DCA-induced suppression of stress-related pathways. Together, these findings highlight metabolic cross-feeding and cooperative stress responses as key features of the *B. thetaiotaomicron*–*C. aerofaciens* interaction under bile acid stress and provide new insights into microbial mechanisms relevant to gut–liver axis physiology.

## Acknowledgements

We thank Kathleen Eismann, Nicole Bock, and Olivia Pleßow for their excellent technical assistance. We also thank Zheng Chen for valuable advice. This work was supported by the Deutsche Forschungsgemeinschaft (DFG, German Research Foundation) – Project-ID 403224013–SFB 1382.

## Declaration of Generative AI Use

The authors used Claude and ChatGPT for language editing and proofreading during manuscript preparation. All scientific content, data analysis, and conclusions were generated and verified by the authors.

## Data Availability Statement

The mass spectrometry proteomics data have been deposited to the ProteomeXchange Consortium via the PRIDE [53] partner repository with the dataset identifier PXD069395.

The metabolome data of all three studies are available at the NIH Common Fund’s National Metabolomics Data Repository (NMDR) website, the Metabolomics Workbench[54], where they have been assigned Study ID ST004535, ST004539 and ST004548. The data can be accessed directly via its Project DOI: http://dx.doi.org/10.21228/M83V8G. This work is supported by NIH grant U2C-DK119886 and OT2-OD030544 grants.

The R scripts used for data processing, statistical analysis, and visualisation are archived on Zenodo at DOI <u>10.5281/zenodo.18297860</u>. The live development version of this code is also available on GitHub at https://github.com/wangyan226/Rscript_BTnCA_DCA.

## Supplement

Supplement_BTnCA_DCA_YW_GM.docx

## Figures and Figure captions

Figures_BTnCA_DCA_YW_GM.docx

## Abbreviations

CA: cholic acid
CDCA: chenodeoxycholic acid
DCA: deoxycholic acid
LCA: lithocholic acid
NAFLD: non-alcoholic fatty liver disease BSH bile salt hydrolase
HSDH: hydroxysteroid dehydrogenase
UDCA: ursodeoxycholic acid
gDCA: glycine-conjugated DCA
IBD: inflammatory bowel disease
IBS: irritable bowel syndrome
*B. theta* / Bt: *Bacteroides thetaiotaomicron*
*C. aero* / Ca: *Collinsella aerofaciens*
qPCR: quantitative PCR
FA: formic acid
PITC: phenylisothiocyanate
KEGG: Kyoto Encyclopedia of Genes and Genomes
OD_600_: optical density at 600 nm
PCA: principal component analysis
BCAA: branched-chain amino acids
VB5: vitamin B5, pantothenate
PLP: pyridoxal 5’-phosphate
SPD: spermidine
SCFA: short-chain fatty acid
ADI: arginine deiminase
QS: quorum-sensing

## Authors’ contributions

YW performed the laboratory experiments, analysed the data, and wrote the manuscript. SBH contributed scripts and coding for data analysis.

AMG contributed the reference genome analysis. ML measured DCA and its derivatives.

DM measured PLP. RP measured PLP. HS measured PLP.

CK supervised the reference genome analysis. URK supervised the metabolomics analyses.

NC supervised the proteomics analyses.

MvB provided overall guidance and edited the manuscript.

## Ethics approval and consent to participate

Not applicable

## Consent for publication

Not applicable

---

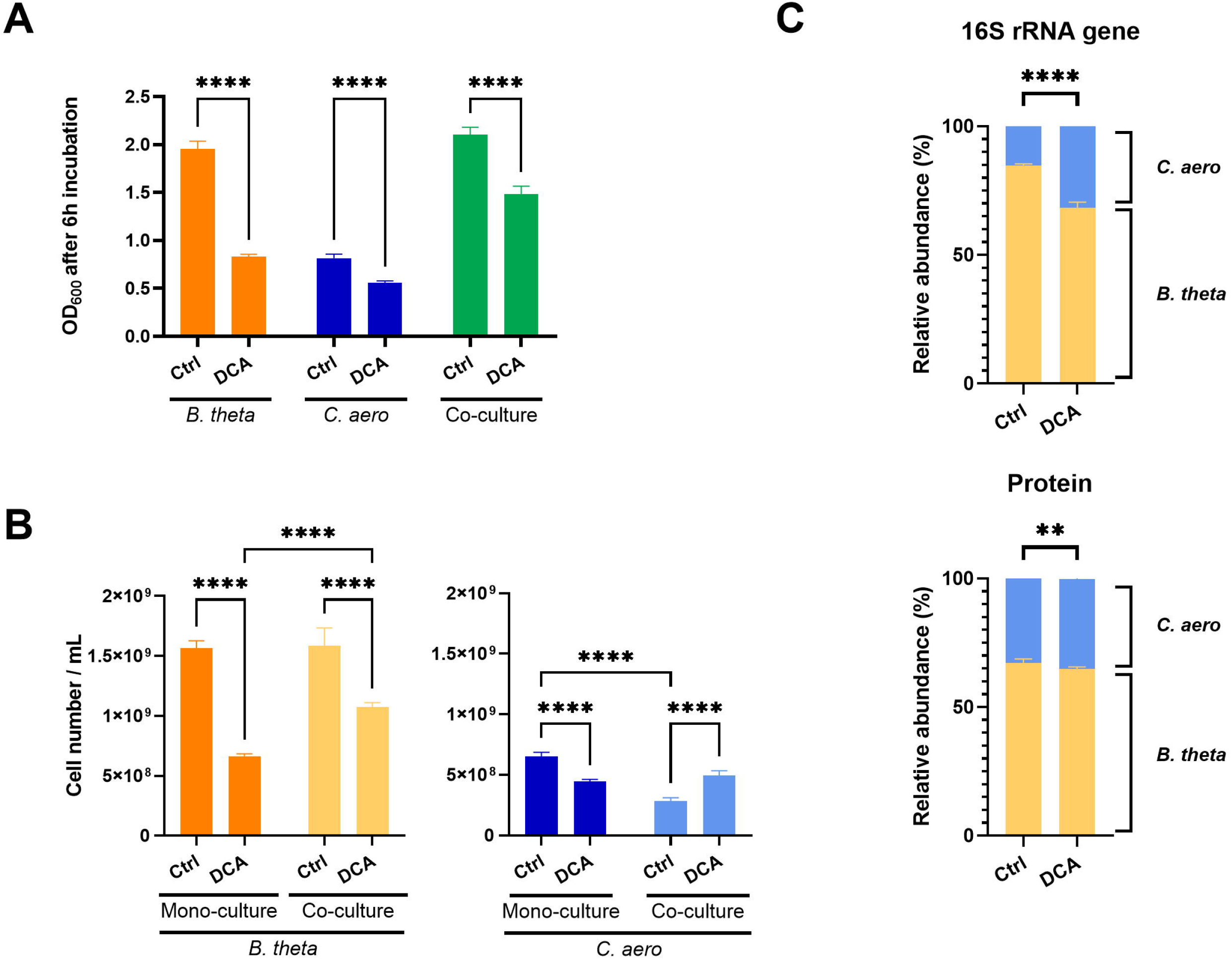

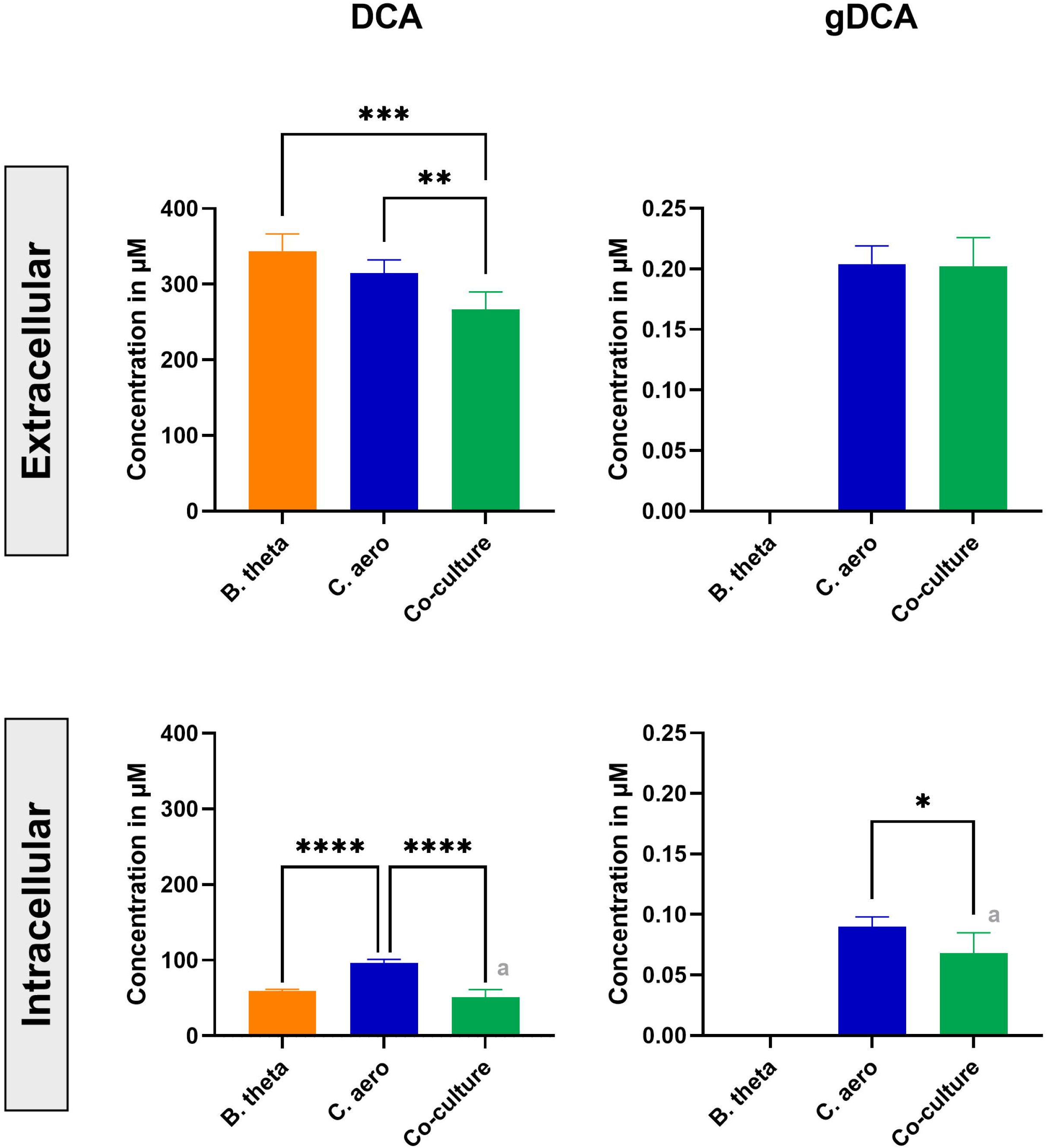

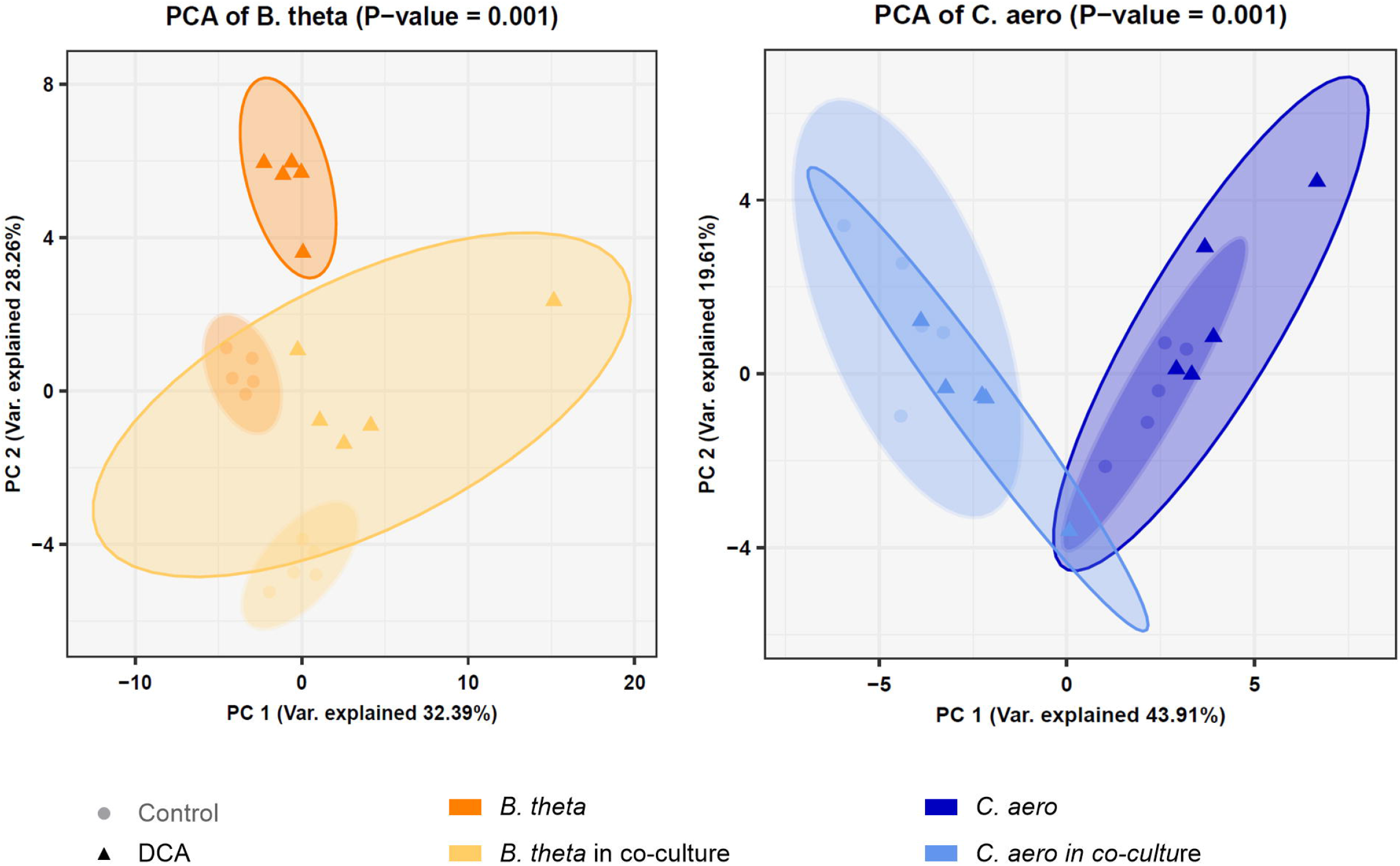

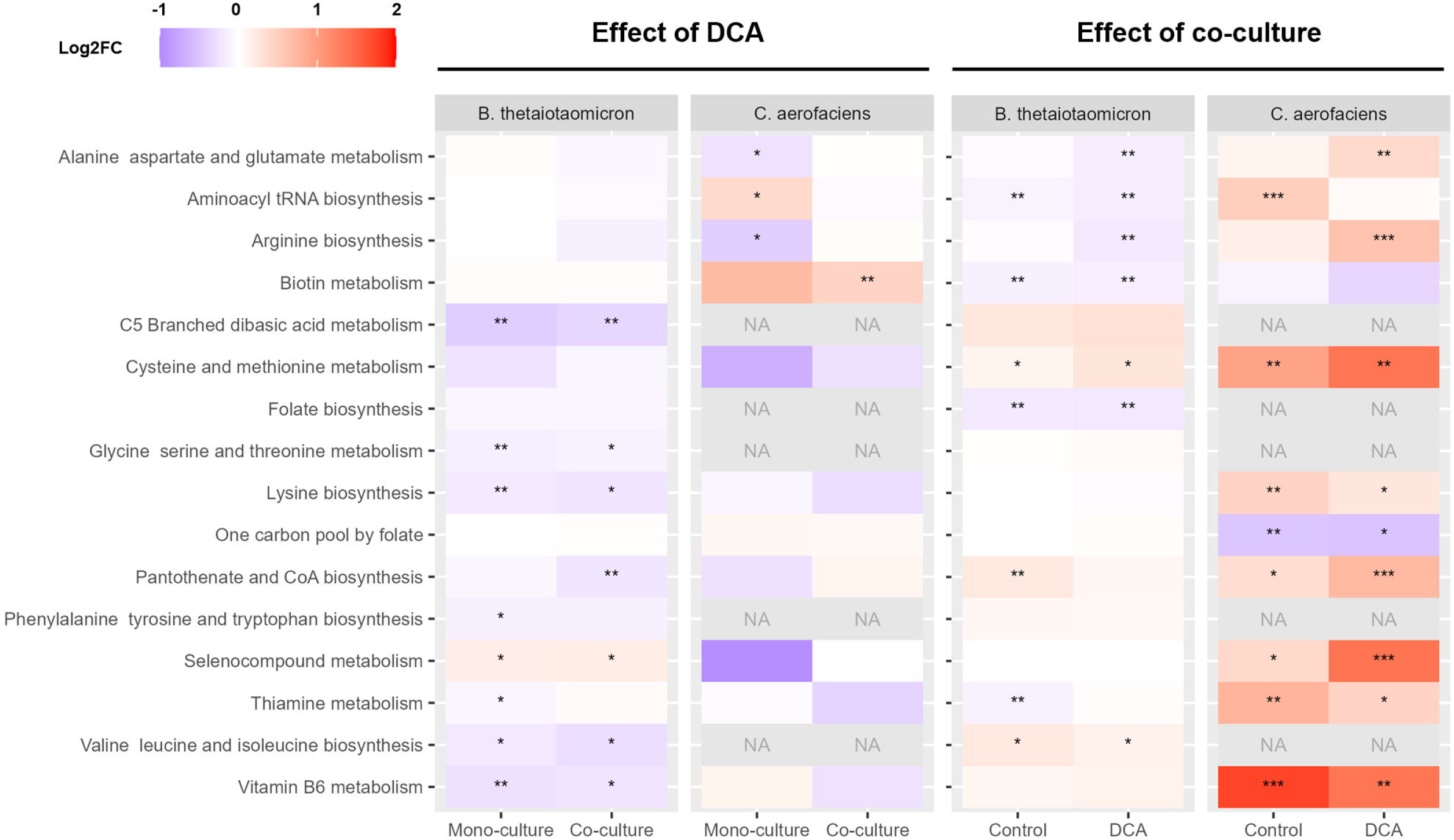

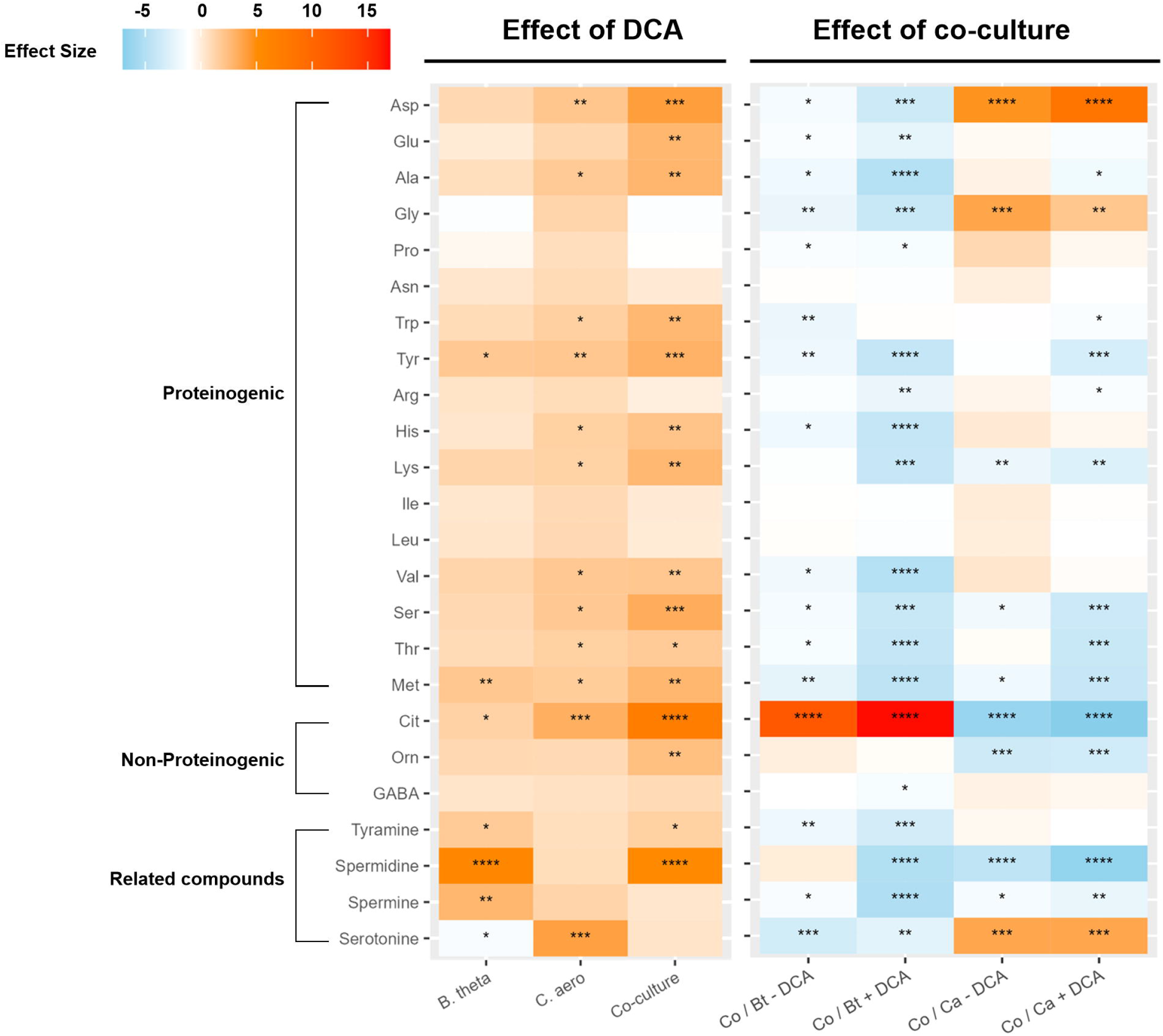

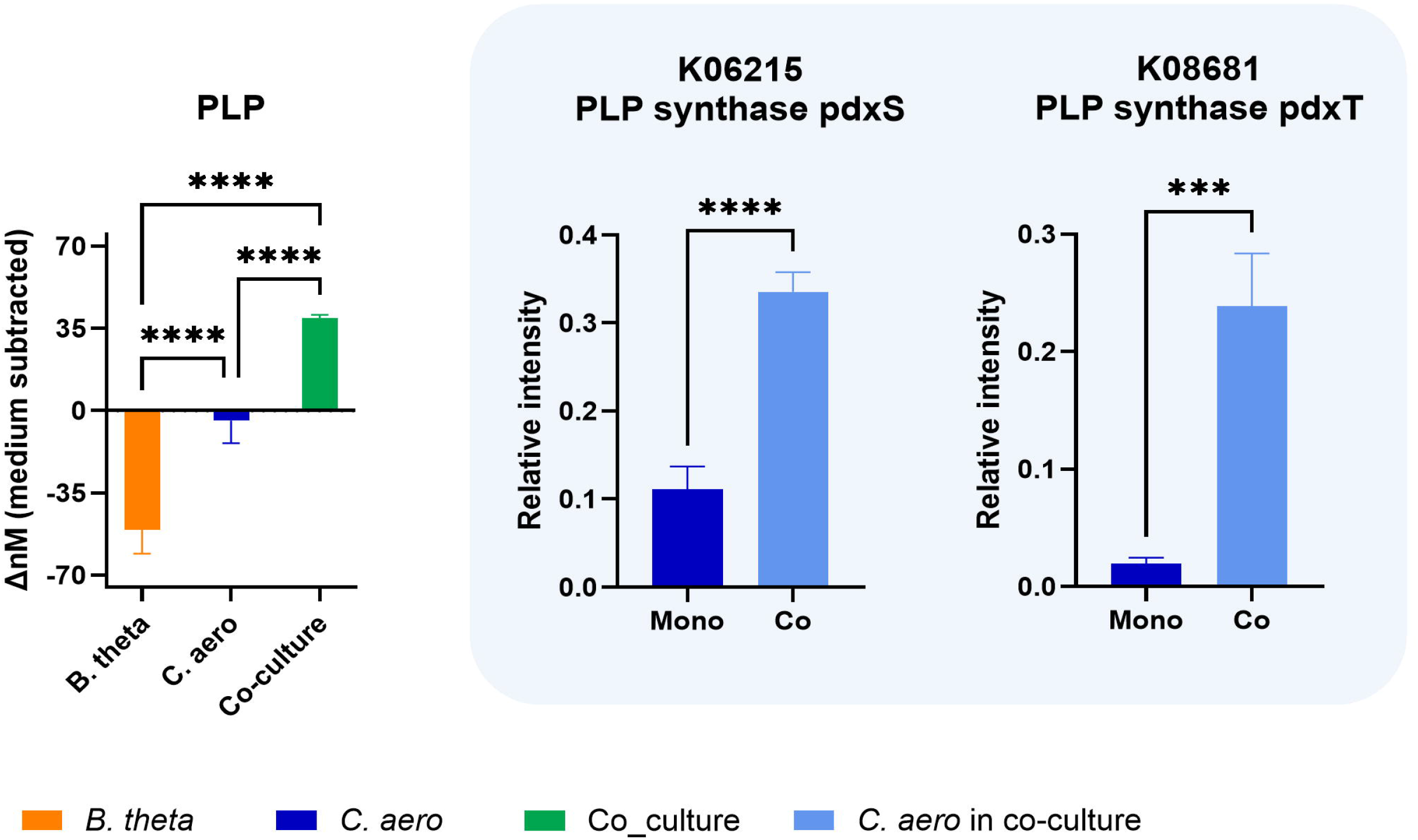

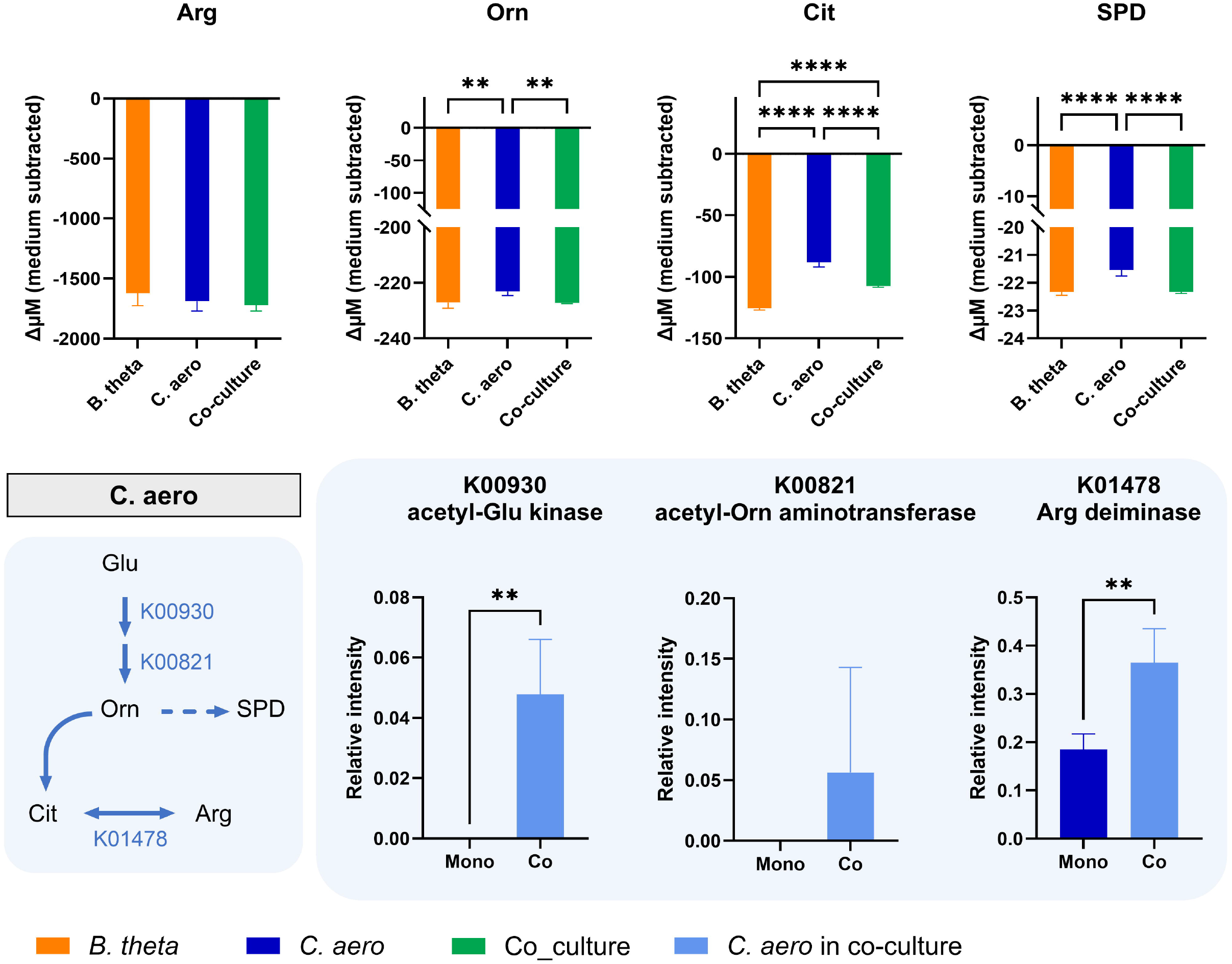

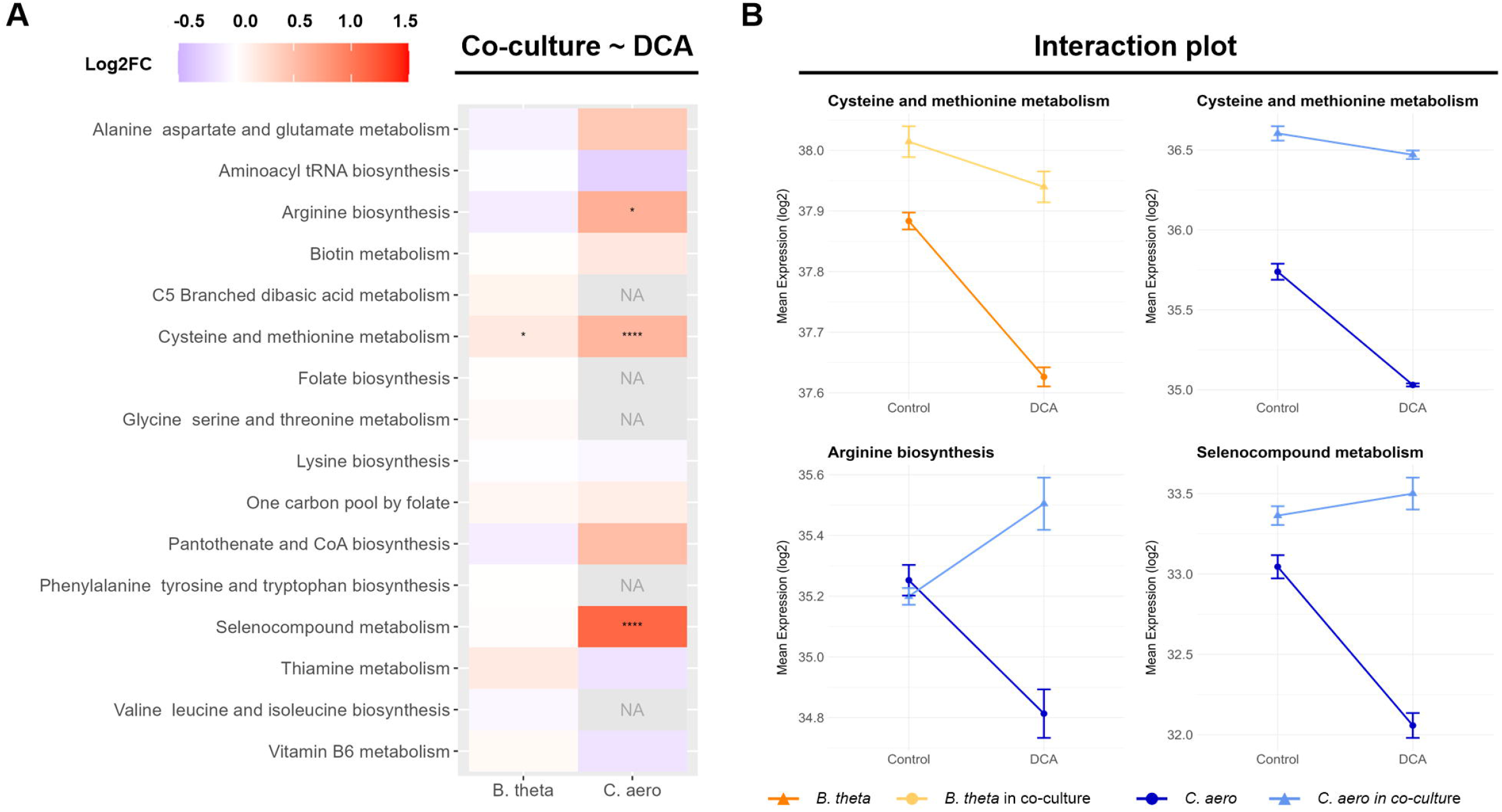

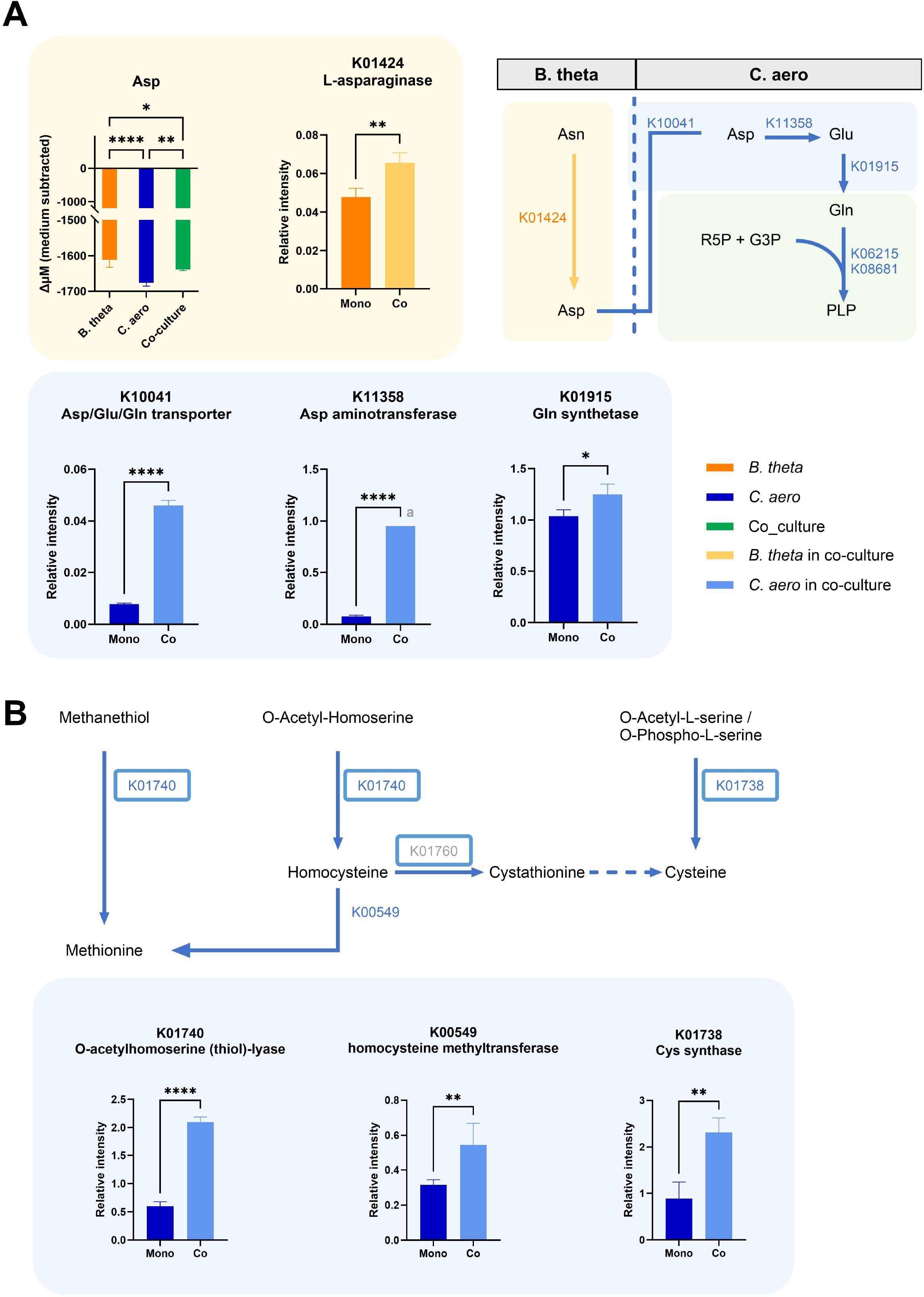

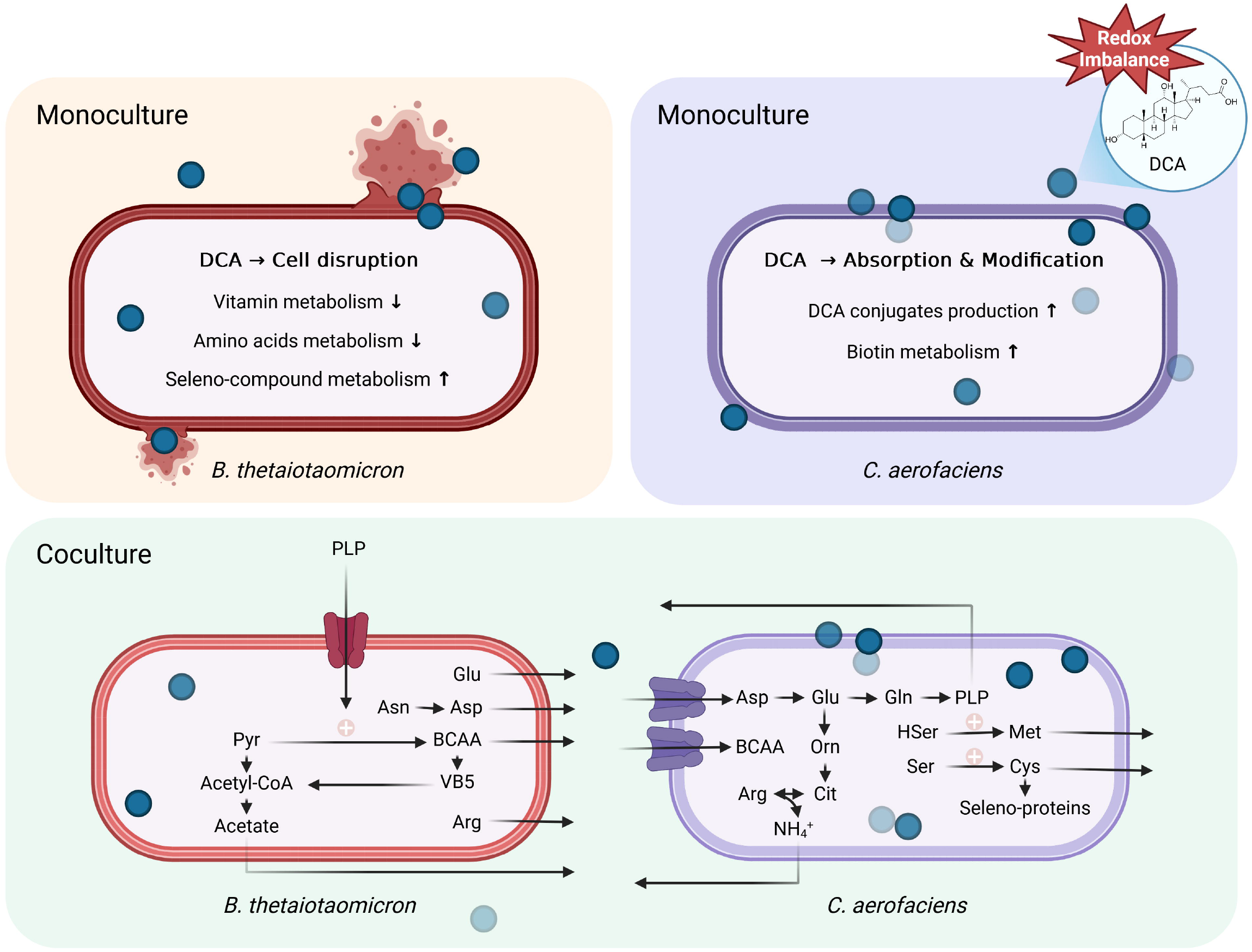

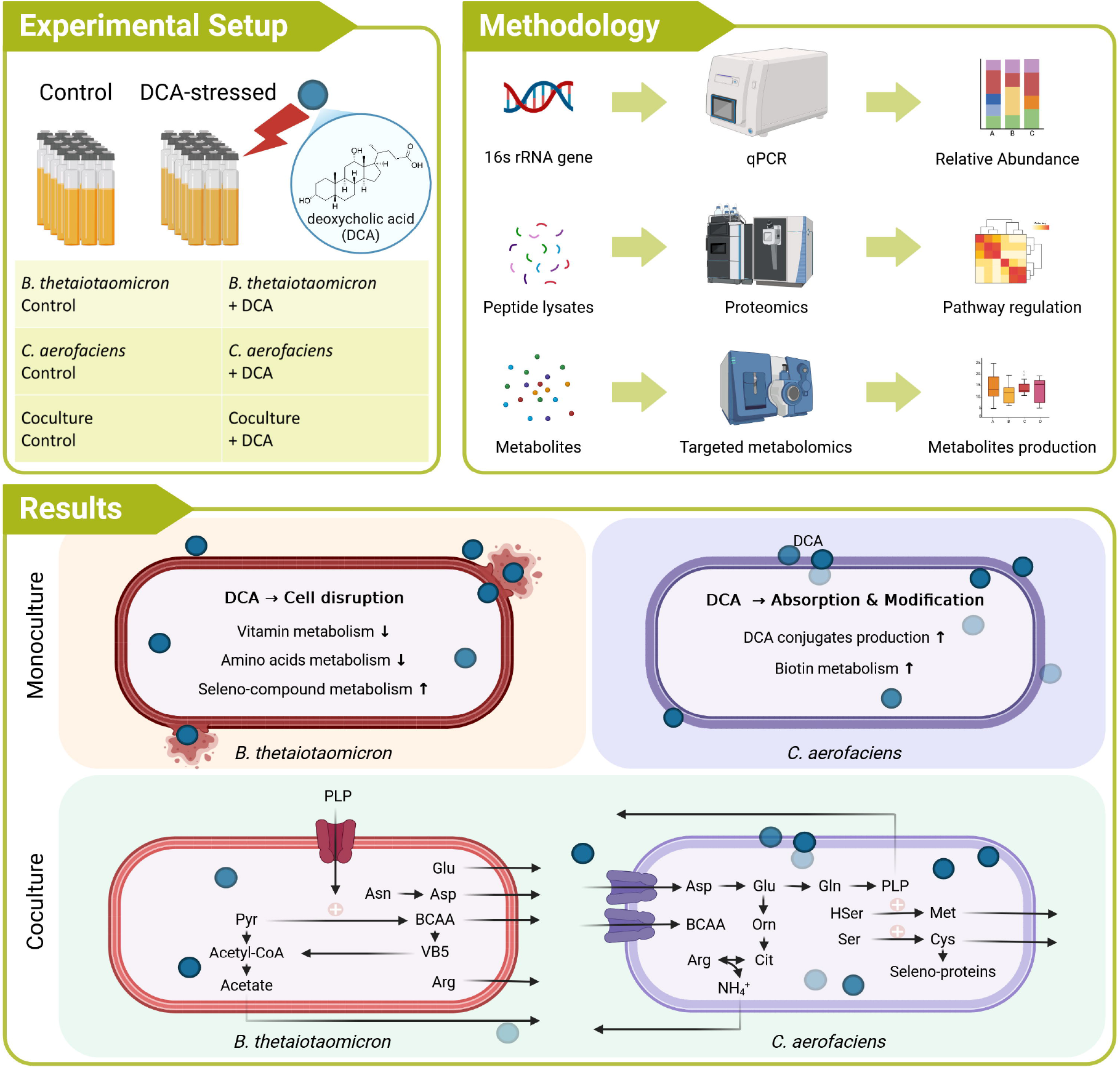

