## Supplement for "Coculture of *Collinsella aerofaciens* and *Bacteroides thetaiotaomicron* under bile acid stress reveals vitamin B6 exchange"

### Bacterial Strains and Cultivation

We cocultured *Bacteroides thetaiotaomicron* and *Collinsella aerofaciens* under deoxycholic acid (DCA) stress, alongside untreated controls, to investigate their metabolic interactions. *B. thetaiotaomicron* (DSMZ 2079) and *C. aerofaciens* (DSMZ 3979) were used in this study. In the figures, they are abbreviated as *B. theta /* Bt and *C. aero* / Ca, respectively. Both strains were cultured in Brain Heart Infusion-supplemented (BHIS) medium under anaerobic conditions at 37 °C with shaking at 175 rpm. The composition of the BHIS medium is provided in Table S1*.*

#### Table S1: Brain Heart Infusion-supplemented (BHIS) medium

| Brain heart infusion broth() | 37.0 g |
| --- | --- |
| Yeast extract | 5.0 g |
| L-Cysteine.HCl | 0.5 g |
| 1 M Resazurin solution | 4.0 mL |
| Hemin-VK1 Solution () | 10.0 mL |
| Distilled water | Fill up to 1.0 L |

The medium was then distributed into Hungate tubes and flushed with pure nitrogen for 1 minute. Sterilization was carried out by autoclaving at 121°C for 20 minutes. The sterile medium was stored at 4°C before use.

### Bile Acid Stress Assay

Prior to the experiment, each strain was pre-cultured individually overnight in BHIS. Hungate anaerobic culture tubes containing BHIS medium were supplemented with DCA to a final concentration of 300 µM and flushed with pure nitrogen gas. Strains were then inoculated into the tubes in five replicates each. Cultures were monitored at 600 nm using a Nanocolour® UV/VIS II spectrophotometer (Macherey-Nagel). After 3 and 6 hours of incubation, 2.5 mL of bacterial suspension was harvested. The taken samples were distributed and centrifuged at 4 °C for 10 minutes to separate the supernatant and pellet. Both fractions were immediately frozen at –80 °C for further analysis.

### Quantitative PCR Assay

In order to determine the proportion of each of the two bacteria in the coculture, an absolute quantitative method was adopted.

*B. thetaiotaomicron* and *C. aerofaciens* were incubated separately overnight, then one millilitre of each bacterial culture was taken to measure OD and centrifuged, and pellets were taken to extract DNA, from which we obtained the DNA extraction rate for each of the two strains. The extracted DNA was used as a standard curve to determine the respective cell numbers in the coculture. The qPCR target gene was the 16S rRNA V4 region gene.

A dye-based qPCR assay was used to quantify bacterial cell counts in turbid CIM cultures, with absolute quantification achieved via species-specific standard curves. Reactions were performed using Luna® Universal qPCR Master Mix (New England Biolabs, 2025), which contains SYBR® Green I for real-time detection of double-stranded DNA and Hot Start Taq DNA polymerase for amplification. Each 20 µL reaction consisted of 10 µL master mix, 0.25 µM of each primer, and a variable amount of DNA template. Thermal cycling was carried out on a 7500 Fast Real-Time PCR System (Applied Biosystems) using the “Fast Ramp” program: initial denaturation at 95°C for 20 s, followed by 40 cycles of 95°C for 3 s and primer-template annealing/extension at 55°C (*C. aerofaciens*) or 60°C (*B. thetaiotaomicron*) for 30 s, with a single fluorescence reading at the end of each cycle. The Luna® mix allows combined annealing and extension in a single step. A continuous melting curve from 60°C to 95°C was recorded after amplification to verify product specificity.

#### Table S2. Primer sequences for absolute quantitative polymerase chain reaction.

| Bacterium | Forward primer (5’ – 3’) | Reverse primer (5’ – 3’) |
| --- | --- | --- |
| *C. aerofaciens* DSM 3979 | TGCTACAATGGCCGGTACAG | AGCAACTCCGACTTCATGG |
| *B. thetaiotaomicron* DSM 2079 | GCAAACTGGAGATGGCGA | AAGGTTTGGTGAGCCGTTA |

#### Chelex DNA Extraction

**Solutions needed**

- 100% Ethanol
- 100% Isopropanol
- 3 M Sodium acetate: ~50 µl per sample
- 7.5 M Ammonium acetate: ~150 µl per sample
- 10% (w/v) Chelex: ~300 µl per sample

**DNA extraction**

- Take 1 ml of bacterial culture
- Centrifuge at 3000 g for 10 min at 4 °C
- Resuspend the pellet with 300 µl Chelex
- Thermomix at 95 °C and 1000 rpm for 50 min
- Centrifuge at 21460 g (R: 128), 3 min
- Transfer the supernatant into a fresh Eppi

**DNA purification (Samples II)**

- + 150 µl NH_4_Ac (final concentration of 2.5 M)
- Incubate for 5 min at -20 °C
- Vortex briefly and centrifuge at 12000 g for 10 min
- Transfer the supernatant into a fresh Eppi (2 ml)
- + 50 µl ice-cold NaAc (final concentration of 0.3 M)
- + 1 ml ice-cold 100% ethanol
- Vortex briefly and incubate at -80 °C for 1.5 h (45 min sufficient)
- Centrifuge at 15000 g for 1 h at 4 °C
- Discard the supernatant
- Wash the pellet with 200 µl ice-cold 100% ethanol
- Centrifuge at 15000 g and 4 °C for 10 min
- Repeat the washing step
- Precipitate with 200 µl ice-cold 100% isopropanol
- Centrifuge at 15000 g and 4 °C for 10 min
- Dry the DNA precipitate to complete dryness (lay laterally on paper, RT for 10 min)
- Solubilise the DNA pellet in 200 µl ddH_2_O

### Metabolite analysis

Intracellular DCA and derivatives were extracted from cell pellets kept on ice. Pellets were resuspended in 500 µL of pre-cooled (–20 °C) methanol and subjected to three freeze–thaw cycles in liquid nitrogen. To ensure cell disruption, suspensions were sonicated for 10 min. After centrifugation at 16,000 g for 5 min at 4 °C, the supernatant was transferred to a fresh tube. The pellet was re-extracted using the same procedure, and both supernatants were combined and dried completely in an Eppendorf® Concentrator Plus at room temperature. Dried extracts were stored at –80 °C until analysis. The supernatant was collected and stored at –80 °C until bile acid analysis. The dried residue was reconstituted in 300 µL of methanol. For DCA quantification, 3 µL of this solution was analysed, while 250 µL was used for the quantification of glyco-, tauro-, and keto-derivatives of DCA. Each aliquot was spiked with an internal standard mixture (d4-DCA, d4-UDCA, d4-gCDCA), evaporated to dryness, and reconstituted in 100 µL of mobile phase. Subsequently, 15 µL were analysed by LC–MS/MS. The sample was separated on a HPLC system (Dionex Ultimate 3000, Dionex Softron GmbH, Germany) equipped with a Pinnacle DB C18 column (100 × 2.1 mm, 3 μm, Restek, USA) and an appropriate guard column. The mobile phase consisted of water, methanol, ammonium acetate, and formic acid (FA); flow rate was 0.3 mL/min, with the column chamber set to 55 °C. Ammonium acetate and FA concentrations at all times were kept at 0.005 M and 0.012% (v/v), respectively, with methanol concentrations (v/v) as follows: 0–2.5 min 40%; 2.5–3.5 min 40–57%; 3.5–9.5 min 57–59%; 9.5–10.0 min 59–70%; 10.0–14.0 min 70–72%; 14.0–16.0 min 72–76%. Then, the column was washed with 95% methanol for 9 min and equilibrated with 40% methanol for 5 min (ammonium acetate and FA were present in both steps). Qualitative reference standards for DCA keto-derivatives were generated using recombinant 3α- or 12α-hydroxysteroid dehydrogenase.

For extracellular amino acid analysis, 10 µL of the culture supernatant was completely dried. The residue was derivatised with 50 µL of phenylisothiocyanate (PITC) solution (5% PITC in ethanol/water/pyridine, 1:1:1, v/v/v) for 25 min at room temperature, then dried again. The resulting residues were resuspended in 10 µL of extraction solution (5 mM ammonium acetate in methanol) by shaking at 1400 rpm for 10 min in a thermomixer (Eppendorf, Hamburg, Germany), and subsequently diluted with a 1:1 mixture of running buffers A and B prior to analysis. Following extraction, analytical samples were first separated using a Waters Acquity™ Ultra-Performance LC system (Framingham, USA) and subsequently analysed and quantified with a QTRAP® 5500 mass spectrometer (SCIEX, Framingham, USA). Chromatographic separation was performed on an Agilent Zorbax Eclipse XDB-C18 column (3.5 µm, 3.0 × 100 mm) at a constant flow rate of 0.5 mL/min at 50 °C with a pre-column from Security Guard Cartridge Kit (Phenomenex, Torrance, USA). The mobile phases consisted of 0.2% FA in water (A) and 0.2% FA in acetonitrile (B). The linear LC gradient was programmed as follows: 0–0.5 min, 0% B; 0.5–4 min, 0–70% B; 4–5.3 min, 70% B; 5.3–5.4 min, 70–0% B; and 5.4–7.3 min, 0% B. The QTRAP was operated in positive ionisation mode. Peak areas of all samples and calibration standards were quantified using SciexOS® software (v3.0.0, SCIEX).

While DCA derivatives and amino acids were analysed using metabolomic approaches, extracellular pyridoxal 5’-phosphate was quantified using a targeted fluorescence-based method. All measurements were performed using the HPLC kit KC2151 (Immundiagnostik AG, Bensheim, Germany). For sample preparation, 100 µL of the sample was mixed with 300 µL of precipitation reagent and centrifuged for 5 min at 10,000 × g. Subsequently, 150 µL of the supernatant was combined with 250 µL of derivatisation solution and incubated for 5 min at 60 °C. After derivatisation, 50 µL of the reaction mixture was injected into the HPLC system. High-performance liquid chromatography (HPLC) analysis was performed using a VWR Hitachi Chromaster Plus HPLC system (Hitachi, Japan) equipped with Clarity VA Chromatography Software (version 8.7.0.94). Separation was achieved under reversed-phase conditions using an Avantor ACE C18-PFP column (5 µm, 125 × 4.6 mm). The mobile phase was delivered isocratically at a flow rate of 2.0 mL·min⁻¹, and the column temperature was maintained at 30 °C. Fluorescence detection was carried out with an excitation wavelength of 320 nm and an emission wavelength of 415 nm. The injection volume was 50 µL.

For short-chain fatty acid analysis, 20 µL of cell culture supernatant was mixed with 20 µL acetonitrile, 20 µL 200 mM 3-nitrophenylhydrazine hydrochloride (3-NPH), and 20 µL 120 mM N-(3-dimethylaminopropyl)-N-ethylcarbodiimide and subsequently incubated in the Thermo Mix at 40 °C for 30 min, shaking at 300 rpm. The derivatized samples were then stored at -20 °C until measurement and diluted with 10% acetonitrile.

A 10 µL aliquot of the diluted derivatized solution was injected into an RSLC UltiMate 3000® system (Thermo Fisher Scientific) coupled to a QTRAP 5500® mass spectrometer (AB Sciex, Framingham, MA, USA). SCFAs were separated using an Acquity UPLC BEH C18 column (1.7 µm; Waters, Eschborn, Germany) with water (0.01% formic acid) and acetonitrile (0.01% formic acid) as mobile phases. The flow rate was set at 0.35 mL/min, with the column maintained at 40 °C. The elution gradient was: 2 minutes at 15% B, a 15-minute gradient from 15% to 50% B, 1 minute at 100% B, and 3 minutes re-equilibration at 15% B. SCFAs were identified and quantified using a scheduled MRM method with specific transitions for each SCFA. Peak areas were analysed using Analyst® Software (v1.6.2, AB Sciex), and the data were exported.

### Metaproteome analysis

For metaproteome analysis, pellet lysis, sample preparation, and measurement procedures followed the protocol of Castañeda-Monsalve et al., 2024. Preserved pellets were resuspended in a lysis buffer containing 8 M urea, 2 M thiourea, and 1 mM phenylmethylsulfonyl fluoride. Bacterial cells were disrupted by bead beating using a FastPrep-24 system (MP Biomedicals, Santa Ana, USA; 5.5 m/s, 1 min, 3 cycles), followed by centrifugation at 10,000 g for 10 min. Vivacon 500 columns equipped with a 10 kDa molecular weight cutoff membrane (Sartorius, Göttingen, Germany) were equilibrated with UA buffer (8M urea in 0.1M Tris/HCl at pH 8.5). and centrifuged at 14,000 g for 20 min at room temperature; all subsequent centrifugation steps were performed under the same conditions unless otherwise specified. A total of 50 µg of protein was loaded onto each column and centrifuged at 14,000 g for 40 min at room temperature. Proteins retained on the filter were incubated with 200 µL of 10 mM dithiothreitol in UA buffer at 37 °C and 600 rpm for 1 min, followed by static incubation for 30 min and centrifugation. The proteins were then treated with 200 µL of 50 mM iodoacetamide in UA buffer at 37 °C and 600 rpm for 1 min, followed by static incubation for 10 min in the dark and centrifugation. Columns were washed twice with 100 µL UB buffer (8 M urea in 0.1 M Tris/HCl, pH 8.0) and centrifuged. Digestion was performed by adding 10 µL UB buffer, 1 µg Trypsin-LysC, and 90 µL of 10 mM ammonium bicarbonate solution, mixing at 600 rpm for 1 min, sealing with parafilm, and incubating overnight. The digested proteins were centrifuged at 14,000 g for 60 min, followed by the addition of 50 µL of 500 mM NaCl and centrifugation. Subsequently, 10 µL of 10% FA was added before desalting. Peptide clean-up was performed using a SOLAµ HRP 96-well plate (Thermo Fisher Scientific). The resulting peptide lysates were finally reconstituted in 0.1% FA for mass spectrometric analysis. Data were processed using Proteome Discoverer software (v2.5, Thermo Fisher Scientific) as previously described.

For each nanoLC-MS analysis, 1 µg of peptide lysate was injected into a Vanquish Neo nanoHPLC system (Thermo Fisher Scientific). Peptides were initially trapped on a C18 reverse-phase trapping column (Acclaim PepMap™ 100, 75 µm × 2 cm, 3 µm particle size, nanoViper, Thermo Fisher Scientific) and subsequently separated on a C18 reverse-phase analytical column (Double nanoViper™ PepMap™ Neo, 75 µm × 150 mm, 2 µm particle size, Thermo Fisher Scientific). Separation was performed using a two-step gradient with mobile phase A (0.01% FA in H₂O) and mobile phase B (80% acetonitrile in H₂O, 0.01% FA). During the first gradient step (95 min), the proportion of mobile phase B was increased from 4% to 30%, followed by a second step (40 min) in which B increased from 30% to 55%. The flow rate was maintained at 300 nL/min. Eluted peptides were ionised via a Nanospray Flex™ Ion Source (Thermo Fisher Scientific) and detected using an Orbitrap Exploris™ 480 mass spectrometer (Thermo Fisher Scientific). Mass spectrometer settings for MS scans included a range of 350–1,550 m/z, resolution of 120,000, an AGC target of 3,000,000 charges, and a maximum injection time of 100 ms. An intensity threshold of 8,000 ions and a dynamic exclusion of 20 s were applied. The top 10 most intense ions were selected for MS/MS analysis using an isolation window of 1.4 m/z, resolution of 15,000, AGC target of 200,000 ions, and maximum injection time of 100 ms.

### Data analysis

Proteomics data quality control, statistical analyses and correlations were performed using in-house written or custom scripts in R. Protein functions and pathway assignments were performed using Ghostkoala in Kyoto Encyclopedia of Genes and Genomes (KEGG). The molecular functions, represented as functional orthologs of *B. thetaiotaomicron* and *C. aerofaciens*, were analysed separately using the KEGG Orthology database. In the figures, they are abbreviated as B. theta / Bt and C. aero / Ca, respectively. Only pathways containing a minimum of 5 proteins and a minimum total coverage of 15% were selected for further analysis. To assess significant changes in species abundance and compound concentration, unpaired Student's *t*-tests were performed in GraphPad Prism 10. Amino acid concentrations were baseline-corrected by subtracting values from fresh medium. Effect sizes of extracellular amino acids were calculated using Hedges’ g, and statistical significance was determined using Student’s t-test. Single-effect analyses of coculture and DCA treatment on *B. thetaiotaomicron* and *C. aerofaciens* were performed by the Kruskal–Wallis test. Dual-effect analysis of coculture and DCA treatment on *B. thetaiotaomicron* and *C. aerofaciens* was performed using two-way ANOVA (limma R package). Adjusted p-values were converted into significance levels using a symbolic representation: p ≤ 0.0001 was indicated as “****”, 0.0001 < p ≤ 0.001 as “***”, 0.01 < p ≤ 0.05 as “**”, and p > 0.05 was considered not significant. These annotations were used for visualising and reporting statistical significance in figures.

### Supplemented Figures

#### Figure S1: Growth curves of *B. thetaiotaomicron* and *C. aerofaciens* monitored by optical density at 600 nm.

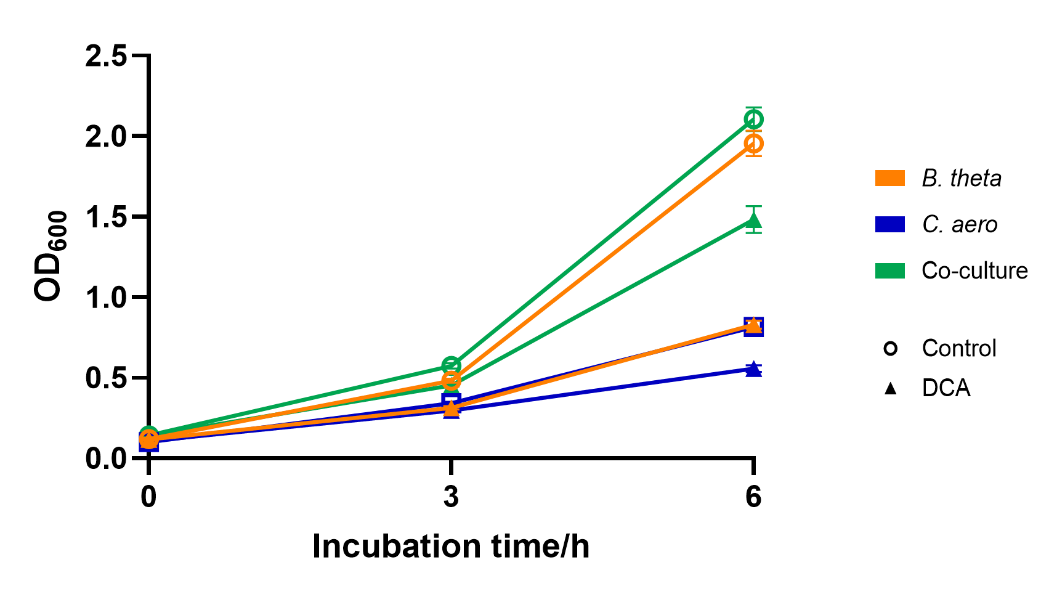

#### Figure S2: Extracellular and intracellular deoxycholic acid (DCA) and its derivatives.

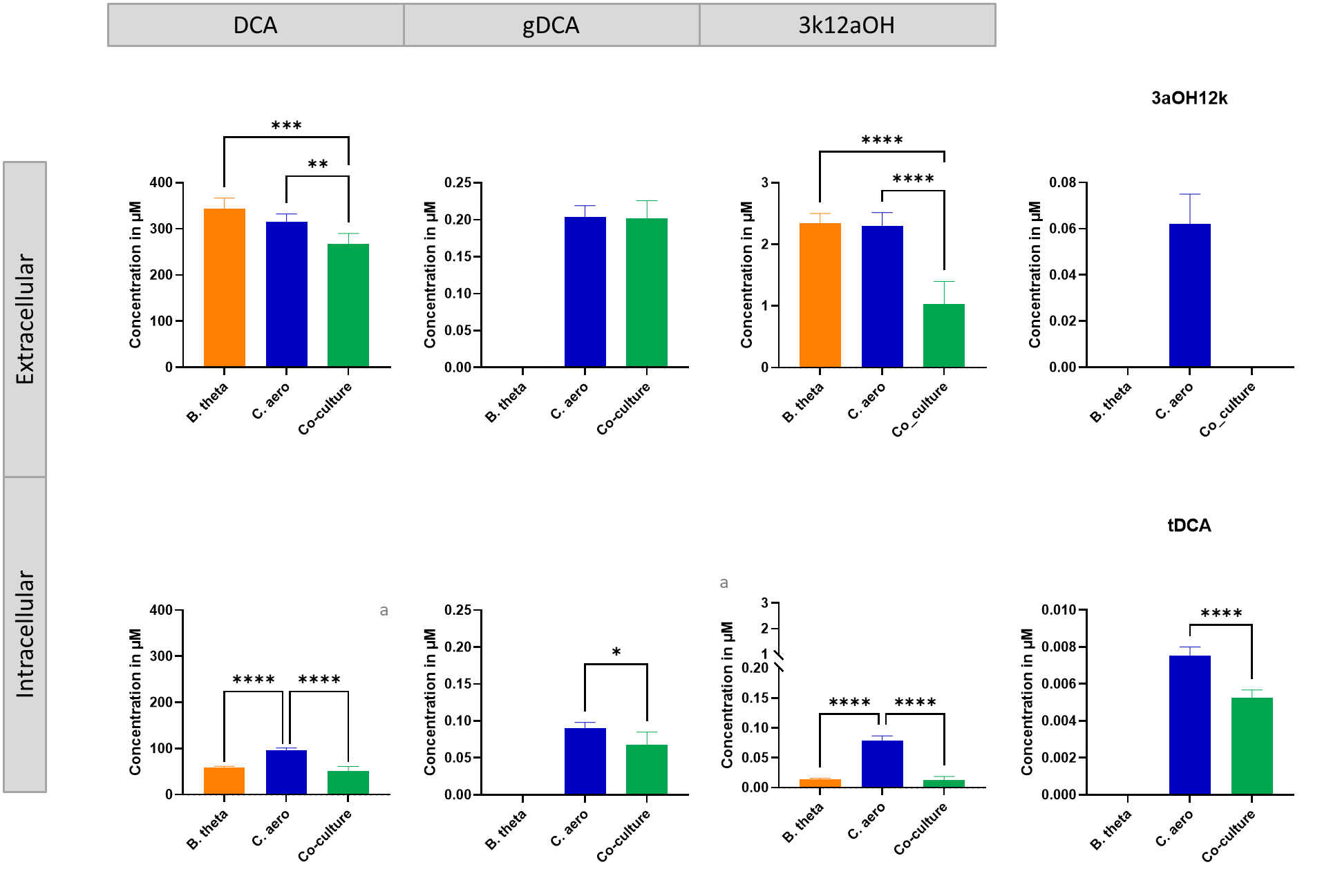

#### Figure S3: Effects of pyridoxal 5’-phosphate (PLP) supplementation on *B. thetaiotaomicron* growth across a concentration gradient.

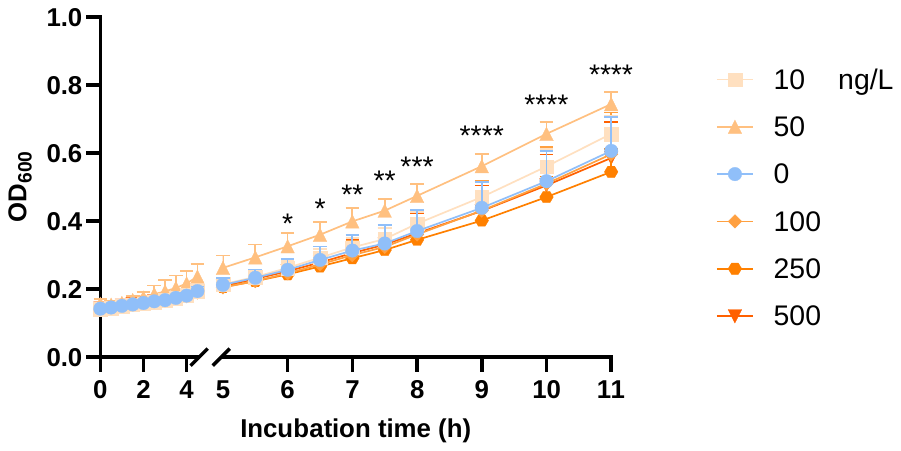

#### Figure S4: Upregulation of enzymes in co-culture potentially linked to increased PLP availability.

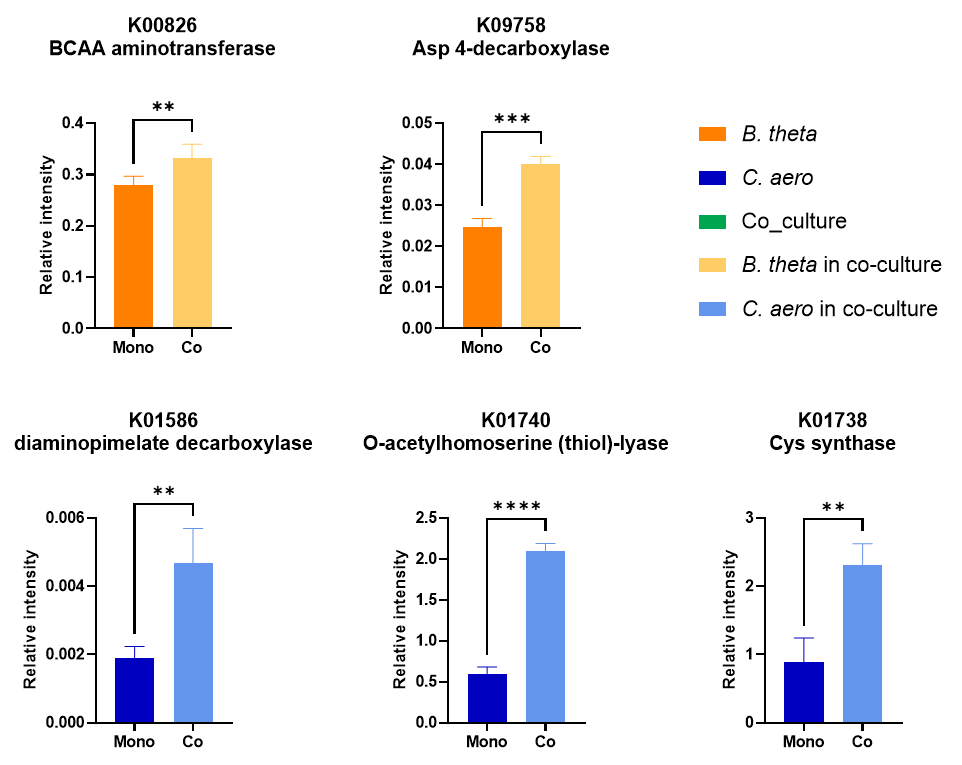

#### Figure S5. Interpretation of branched-chain amino acid metabolism using valine as an example.

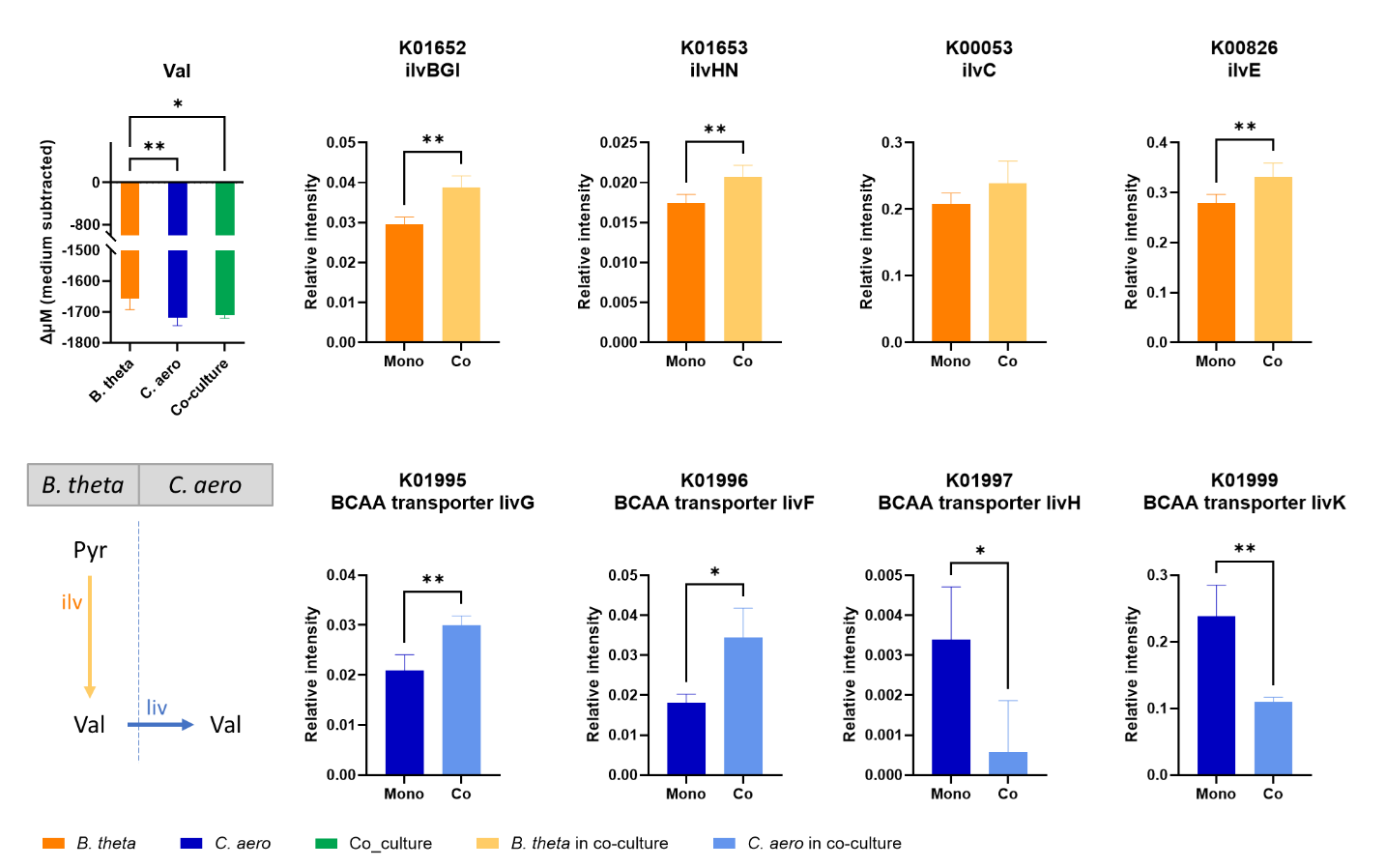

Based on intracellular proteomics and extracellular metabolomics. Data were analyzed and visualized using GraphPad Prism.

#### Figure S6. Extracellular short-chain fatty acids (SCFA)

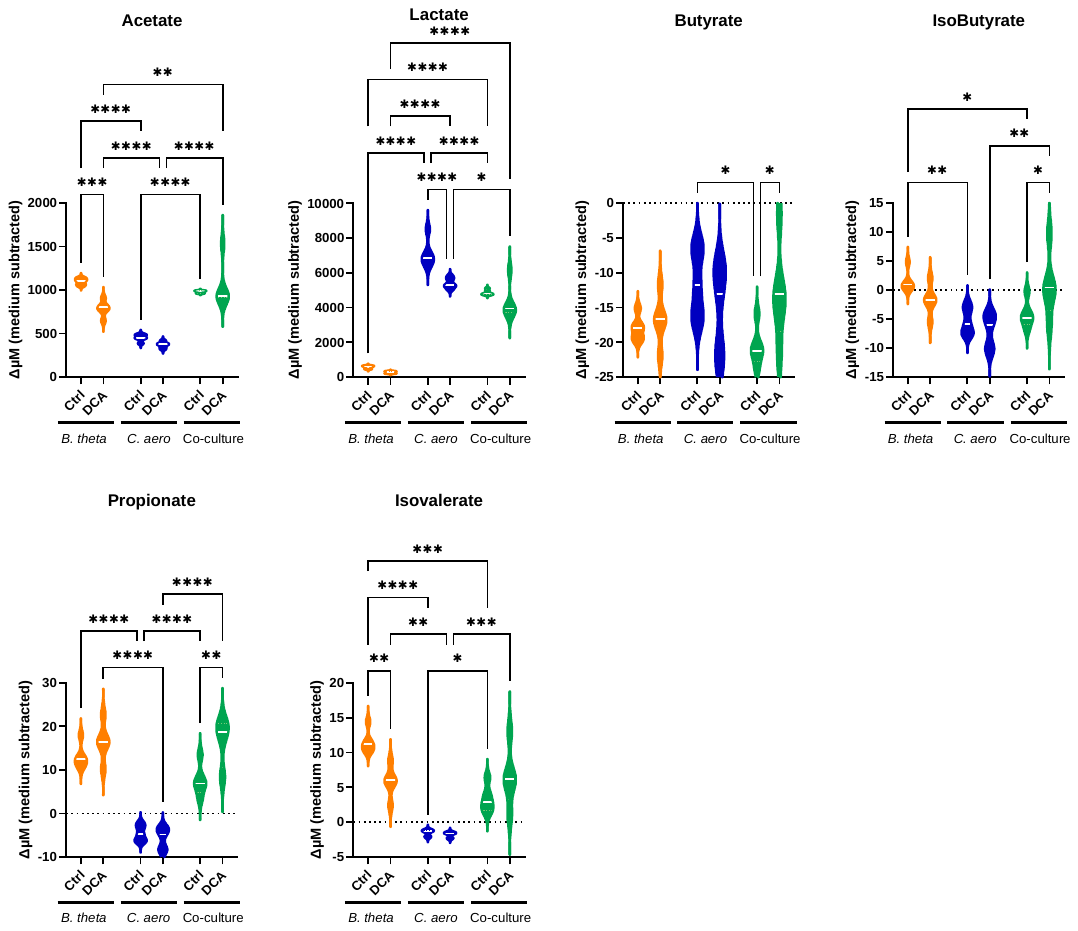

#### Figure S7. Complete *B. thetaiotaomicron* peptide abundance alteration without pathway filtration

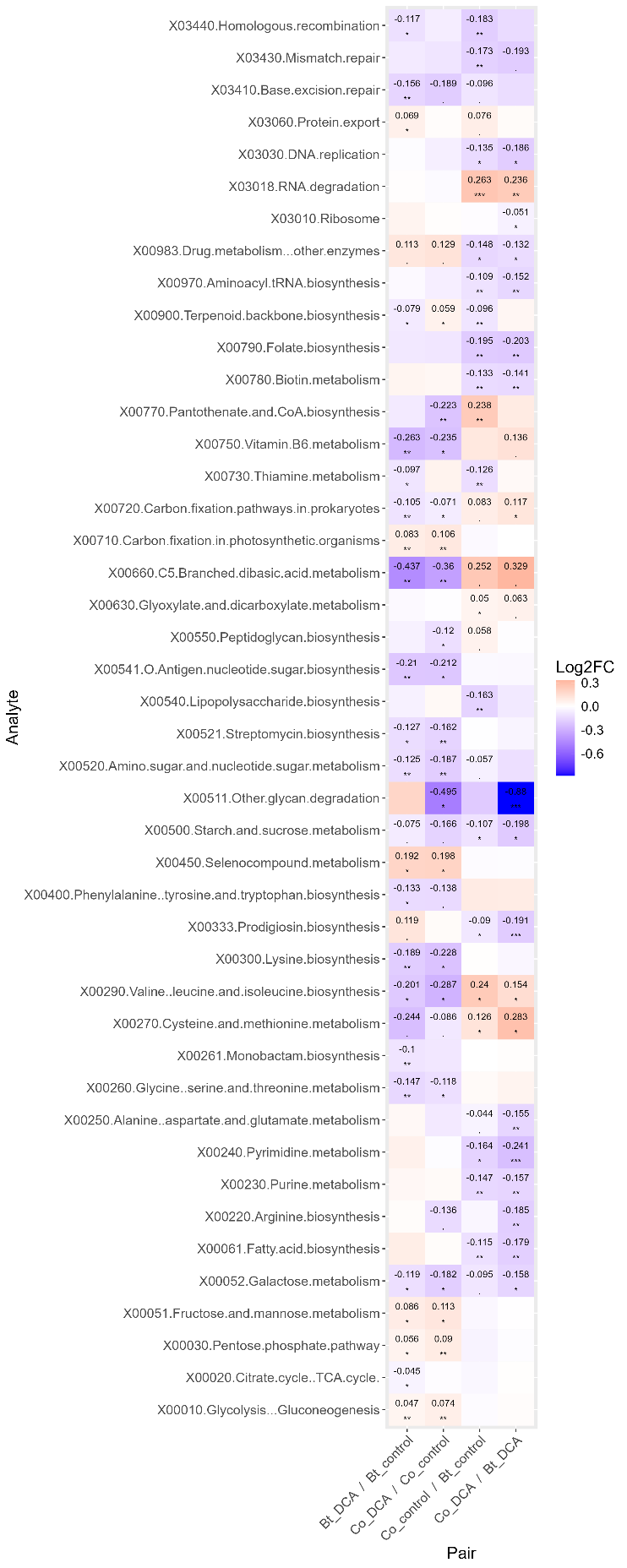

#### Figure S8. Complete *C. aerofaciens* peptide abundance alteration without pathway filtration

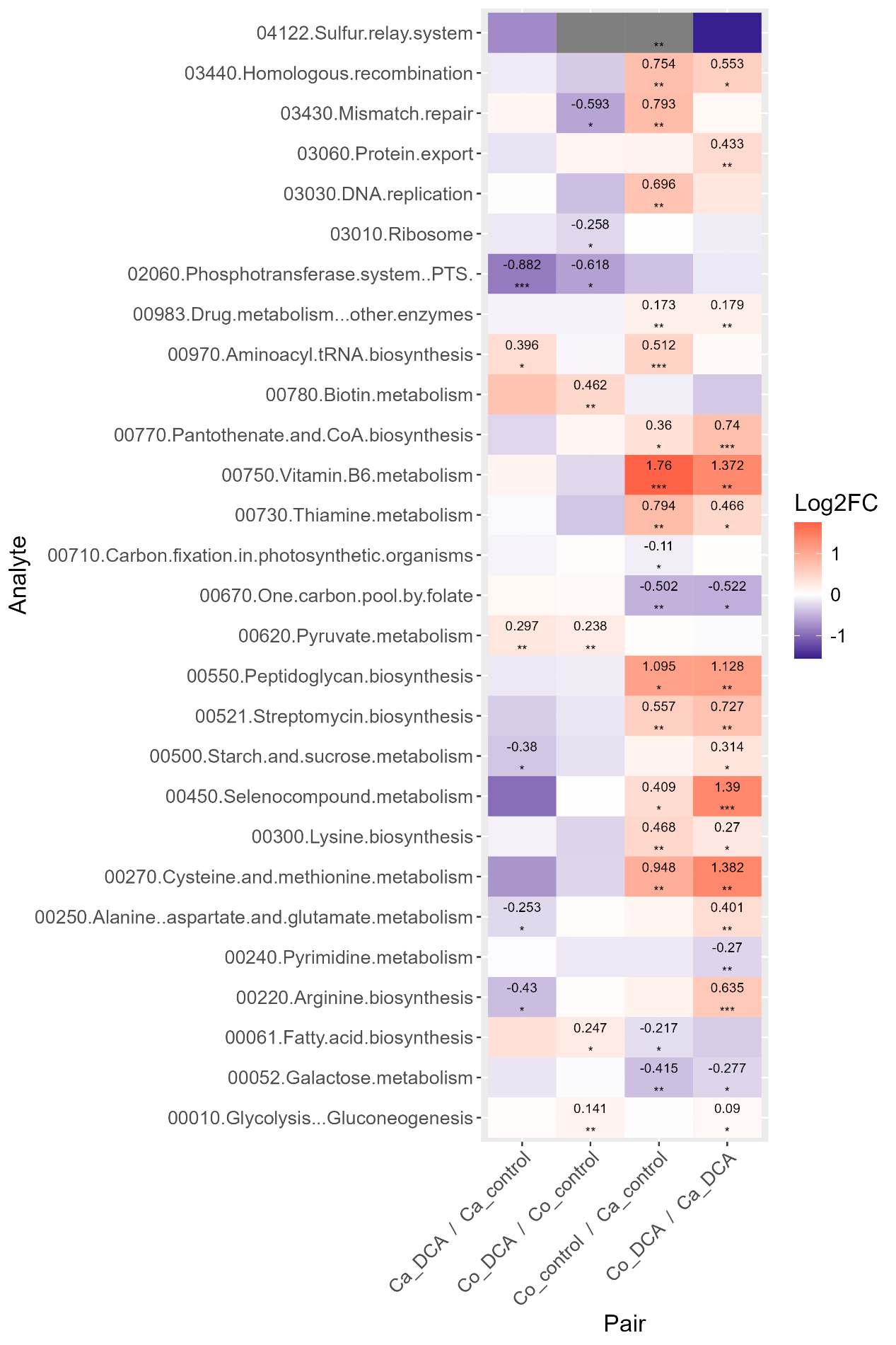
